# Endothelial Cell-Specific Molecule-1 Inhibits Albuminuria in Diabetic Mice

**DOI:** 10.1101/2021.10.14.464296

**Authors:** Xiaoyi Zheng, Lauren Higdon, Alexandre Gaudet, Manav Shah, Angela Balistieri, Catherine Li, Patricia Nadai, Latha Palaniappan, Xiaoping Yang, Briana Santo, Brandon Ginley, Xiaoxin X. Wang, Komuraiah Myakala, Pratima Nallagatla, Moshe Levi, Pinaki Sarder, Avi Rosenberg, Jonathan S. Maltzman, Nathalie de Freitas Caires, Vivek Bhalla

## Abstract

Diabetic kidney disease (DKD) is the most common cause of kidney failure in the world, and novel predictive biomarkers and molecular mechanisms of disease are needed. Endothelial cell-specific molecule-1 (Esm-1) is a secreted proteoglycan that attenuates inflammation. We previously identified that a glomerular deficiency of Esm-1 associates with more pronounced albuminuria and glomerular inflammation in DKD-susceptible relative to DKD-resistant mice, but its contribution to DKD remains unexplored. In this study, we show that lower circulating Esm-1 predicts progressive stages of albuminuria in patients with diabetes. In DKD-susceptible mice, Esm-1 inversely correlates with albuminuria and glomerular leukocyte infiltration. Using hydrodynamic tail-vein injection, we show that over-expression of either mouse or human Esm-1 reduces diabetes-induced albuminuria relative to saline-injected controls independent of leukocyte infiltration. Using a complementary approach, we find that constitutive deletion of Esm-1 in DKD-resistant mice increases the degree of diabetes-induced albuminuria versus wild-type controls. Mechanistically, over-expression of Esm-1 attenuates diabetes-induced podocyte injury. By glomerular RNAseq, we identify that Esm-1 attenuates diabetes-induced up-regulation of interferon-stimulated genes, and Esm-1 inhibits expression of kidney disease-promoting and interferon-related genes, including *Ackr2* and *Cxcl11*. In conclusion, we demonstrate that Esm-1 protects against diabetes-induced albuminuria, and podocytopathy, possibly through select interferon signaling.

## Introduction

Among the many complications of diabetes mellitus, kidney disease has the highest morbidity and mortality^1-3^. The prevalence of diabetic kidney disease (DKD), defined as macroalbuminuria and/or impaired kidney function, is only 15%; and few demographic or clinical features reliably predict which patients with diabetes (Type 1 or Type 2) will develop DKD^4-7^. A better understanding of the molecular determinants of DKD could yield meaningful implications for human health.

Investigators of the Animal Models of Diabetic Complications Consortium have phenotypically characterized the clinical response to hyperglycemia among genetically distinct mouse strains^8, 9^. One of these strains (DBA/2, DKD-susceptible) develops significant nephropathy compared with the more widely used strain (C57BL/6, DKD-resistant)^10^. Glomerular leukocyte infiltration is also significantly higher in DKD-susceptible vs. DKD-resistant mice^11, 12^. We previously compared glomerular transcriptome profiles from these two strains, and in DKD-susceptible mice, we found significantly higher expression of pro-inflammatory genes^13^ and of genes related to endothelial dysfunction^14^ and to immune-mediated vascular injury^15, 16^. Among differentially regulated genes, we showed a glomerular-specific deficiency of Esm-1 in DKD-susceptible mice, and that Esm-1 acutely inhibits leukocyte infiltration^13, 17^.

Esm-1 is a glomerular-enriched, secreted glycoprotein with putative functions in inflammation and developing vasculature, although its effects on the kidney are unknown. Esm-1 transcription is modulated by pro-inflammatory cytokines including gamma-interferon^18^. One putative function of Esm-1 is to inhibit lymphocyte function-associated antigen-1 (LFA-1): intercellular adhesion molecule-1 (ICAM-1) interaction^17, 19^, a well-recognized step of leukocyte infiltration. During development, Esm-1, regulated by VEGF, is also expressed in tip cells at the leading edge of vascular outgrowths^20^. In samples from healthy volunteers, we showed that Esm-1 is enriched in kidney compared to other tissues and within the kidney, is highest in glomeruli^13^. However, this glomerular-enriched Esm-1 is relatively deficient in patients with DKD vs. healthy volunteers^13, 21,22^. We and others have shown that Esm-1 reduces leukocyte transmigration in vitro^13, 17^, and Esm-1 glomerular mRNA and protein are decreased in DKD-susceptible vs. DKD-resistant mice and demonstrate an attenuated increase with hyperglycemia^13^. In contrast, circulating Esm-1 inversely correlates with estimated GFR^23^. Whether circulating Esm-1 ameliorates DKD or is elevated simply due to diminished renal clearance has not been previously studied.

We hypothesize that Esm-1 deficiency in DKD-susceptible vs. DKD-resistant mice contributes to kidney injury in DKD-susceptible mice. Further, we posit that addition of Esm-1 in DKD-susceptible mice would attenuate kidney injury, and Esm-1 deletion in DKD-resistant mice would promote kidney injury. In the present study, using complementary approaches, we determine the in vivo contribution of circulating Esm-1 to albuminuria and glomerular and/or tubulointerstitial inflammation in DKD. We also analyze the histologic and transcriptomic response to Esm-1 rescue in DKD-susceptible mice.

## Results

### Esm-1 deficiency predicts progressive albuminuria in patients with diabetes

We measured circulating Esm-1 in patients with diabetes with and without albuminuria. Among patients with diabetes, those with micro- or macroalbuminuria demonstrate significantly lower estimated GFR than those with normoalbuminuria (**Table 1, Supplementary Figure 1**). In a cross-sectional analysis, circulating Esm-1 does not inversely correlate with kidney injury in diabetic patients although the level of Esm-1 may be confounded by stage of chronic kidney disease (**Figure 1**). However, in a prospective study, patients with normo- or micro-albuminuria that progressed to micro- or macroalbuminuria have significantly lower baseline Esm-1 (**Figure 1, Table 2**) (Median (Q1-Q3): 5.1 (4.0-5.3) vs 6.8 ng/mL (5.4-9.1), p-value < 0.01). To determine whether these observations extend to our mouse model, we measured circulating Esm-1 in DKD-susceptible mice.

**Table 1:** Baseline characteristics of patients with diabetes mellitus by albuminuria.

|  | **Normoalbuminuria** | **Microalbuminuria** | **Macroalbuminuria** | **p-value** |
| --- | --- | --- | --- | --- |
| **N #** | 54 | 23 | 22 |  |
| **Male N (%)** | 34 (63) | 15 (65.2) | 54.5 (12) | 0.73 |
| **Age (years)** | 64.0 (14.8) | 70.0 (16.7) | 68.0 (20.2) | 0.21 |
| **Hemoglobin A1c (%)** | 6.5 (1.4) | 6.7 (2.1) | 6.9 (1.1) | 0.38 |
| **eGFR (mL/min/1.73m<sup>2</sup>)** | 78.6 (33.6) | 52.8 (22.6)* | 49.7 (30.7)* | <0.01 |
| **Urine ACR (mg/g)** | 21.0 (20.7) | 89 (102.5)* | 1256.5 (1863.6)*# | <0.01 |
| **Total Esm-1 (ng/mL)** | 6.2 (4.0) | 6.3 (3.7) | 5.0 (5.1) | 0.39 |

eGFR, estimated glomerular filtration rate; ACR, albumin-to-creatinine ratio; Data reported as median (interquartile range); \* p-value < 0.05 vs. normoalbuminuria; # p-value < 0.05 vs. microalbuminuria using a Kruskal-Wallis test.

**Figure 1:**
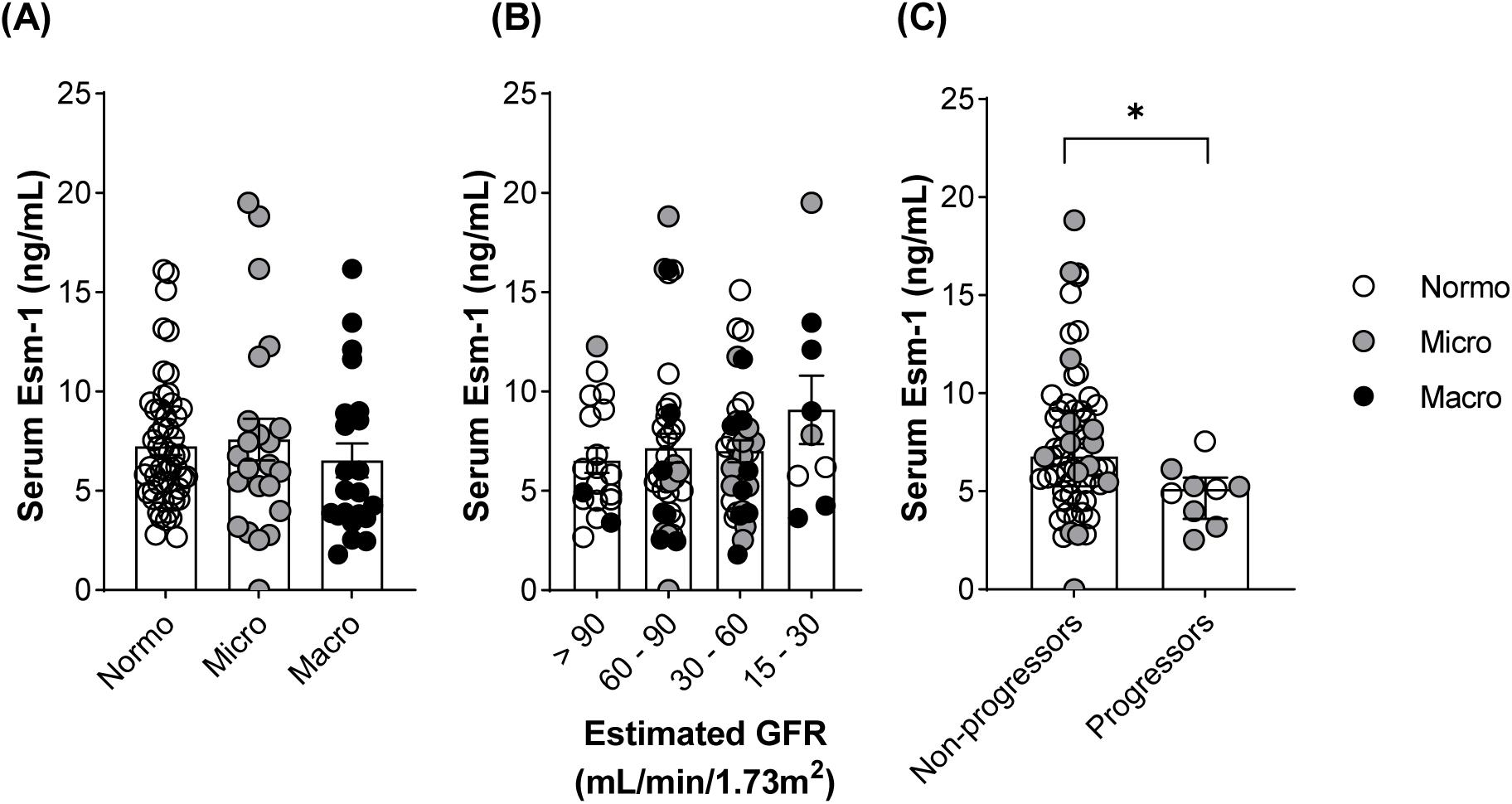
Serum Esm-1 deficiency predicts progressive stages of albuminuria in patients with diabetes. (A) Level of circulating Esm-1 by classification of albuminuria. (B) Correlation of circulating Esm-1 with stages of chronic kidney disease. (C) Level of circulating Esm-1 in non-progressors and progressors. White circles, patients with diabetes with normoalbuminuria; Grey circles, patients with microalbuminuria; and Black circles, patients with macroalbuminuria. Results are presented per patient and include mean ± SEM. *\** p-value < 0.05 vs. non-progressors by unpaired Student’s t-test. N= 22-54 patients per classification of albuminuria.

**Table 2:** Baseline characteristics of patients with diabetes mellitus by progression status.

|  | **Non-Progressors** | **Progressors** | **p-value** |
| --- | --- | --- | --- |
| **N #** | 62 | 9 |  |
| **Male N (%)** | 41 (66.1) | 4 (44.4) | 0.30 |
| **Age (years)** | 64.0 (19.1) | 66.0 (15.9) | 0.28 |
| **Hemoglobin A1c (%)** | 6.7 (1.7) | 7.8 (1.9) | 0.25 |
| **eGFR (mL/min/1.73m<sup>2</sup>)** | 77.1 (31.8) | 54.2 (15.8)* | 0.007 |
| **Urine ACR (mg/g)** | 29.9 (19.8) | 45.0 (70.7)* | 0.01 |
| **Total Esm-1 (ng/mL)** | 6.8 (3.8) | 5.1 (1.3)* | 0.01 |

eGFR, estimated glomerular filtration rate; ACR, albumin-to-creatinine ratio; Data for age, hemoglobin A1c, eGFR, urine ACR, and total Esm-1 are reported as median (interquartile range); \* p-value < 0.05 vs. non-progressors using a Mann-Whitney test.

### Circulating Esm-1 inversely correlates with albuminuria and glomerular leukocyte infiltration in mice with DKD

To establish a model of diabetic kidney disease in mice, we induced diabetes in DKD-susceptible mice with five consecutive doses of streptozotocin (STZ) injection followed by high fat feeding, as indicated in **Figure 2A**. Compared with non-diabetic controls, diabetic DKD-susceptible mice show similar plasma Esm-1 concentrations, but significantly higher albuminuria (**Figure 2B,C**). Overall, albuminuria inversely correlates with plasma Esm-1 concentration (R^2^ = 0.7743), but there are two distinct responses to induction of diabetes. For mice with low plasma Esm-1, diabetes induces significantly higher albuminuria than observed in non-diabetic controls, whereas in similarly treated mice with a plasma Esm-1 > 3 ng/mL, diabetes has virtually no effect on albuminuria (**Figure 2D**).

**Figure 2:**
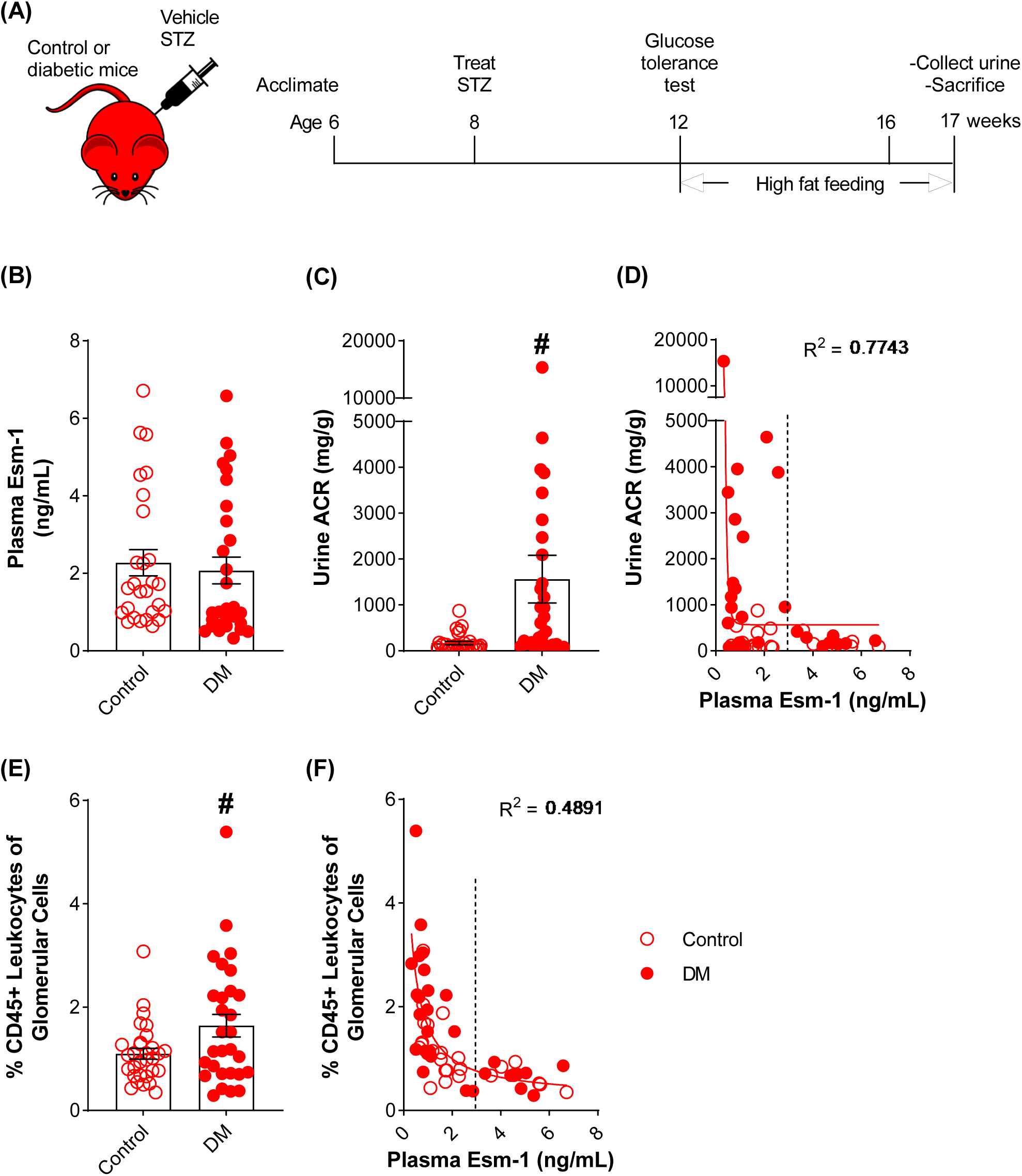
Esm-1 inversely correlates with albuminuria in diabetic, DKD-susceptible mice. (A) Timeline of experimental design. We treated mice with vehicle vs. streptozotocin (STZ) and high fat feeding, followed by sacrifice at 17 weeks of age. We quantified (B) plasma Esm-1 (N= 27-29 per group), and (C) urine albumin-to-creatinine ratio (ACR) (N= 30-31 per group). (D) We correlated urine ACR with plasma Esm-1 (N= 27-29 per group). Dotted line represents plasma Esm-1 of 3 ng/mL. Open circles, control mice; closed circles, diabetic (DM) mice. Results are presented per mouse and include mean ± SEM. R^2^ for non-linear regression (solid line) is shown. # p-value < 0.05 vs. control mice by unpaired Student’s t-test.

We utilize flow cytometry analysis to quantify mouse glomerular leukocyte infiltration. We quantified mouse kidney leukocyte infiltration in both glomeruli and tubulointerstitium (**Supplementary Figure 3A**). We show that 1.1% ± 0.1 and 2.8% ± 0.7 of cells are CD45(+)-positive leukocytes in healthy mice glomeruli and tubulointerstitium (N=29-30 mice per group), respectively, and the predominant leukocyte subtype are macrophages (**Supplementary Figure 3B**,**C**). Diabetic mice have more pronounced glomerular infiltration compared with control mice (**Figure 2E**). Similar to the pattern of albuminuria, plasma Esm-1 inversely correlates with levels of glomerular leukocyte infiltration (R^2^ = 0.49), but for mice with plasma Esm-1 > 3 ng/mL, diabetes has virtually no effect on leukocyte infiltration (**Figure 2F**). The above experiments indicate lower Esm-1 correlates with markers of kidney disease. To assess causality, we utilized over-expression and knockout of Esm-1.

### Hydrodynamic injection stably increases circulating Esm-1 in mice

Intraperitoneally-injected Esm-1 protein only transiently increases plasma Esm-1 (**Supplementary Figure 2**). Therefore, we utilized hydrodynamic tail vein injection. Esm-1 and *Gaussia* luciferase cDNA integrate with a transposase into the genome within hepatocytes for stable over-expression and secretion into the blood (**Figure 3A,B**)^24^. Co-injection of luciferase significantly increases plasma luciferase activity for two weeks. Although plasma luciferase activity plateaus over time, the luciferase activity at both one week and twelve weeks correlates with plasma Esm-1 (**Figure 3C,D**) at those intervals, suggesting plasma luciferase activity is a surrogate marker of Esm-1 over-expression after hydrodynamic injection.

**Figure 3:**
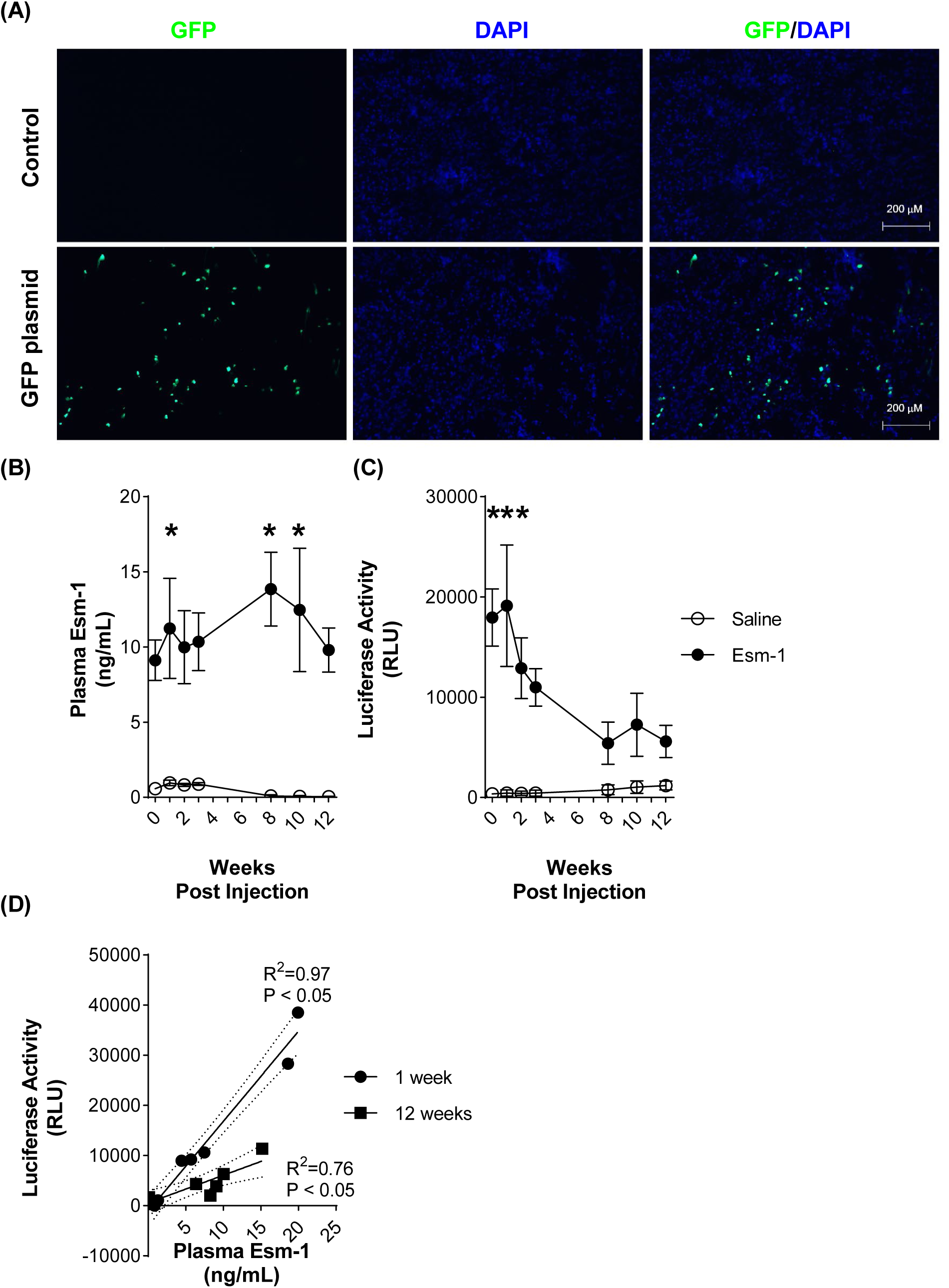
Hydrodynamic injection induces over-expression in hepatocytes and a stable increase in circulating Esm-1. (A) Representative image of GFP fluorescence in liver sections, three days after hydrodynamic tail-vein injection. Scale bars (*white*) as indicated. *Green*, GFP. *Blue*, DAPI. (B) Plasma Esm-1 and (C) luciferase activity in mice treated with saline (control) or Esm-1, Gaussia luciferase, and Sleeping Beauty 100 (SB100) transposase by hydrodynamic injection. Results are presented as mean ± SEM. (D) Correlation of luciferase activity and plasma Esm-1 level at week-1 (circles) and week-12 (squares). Results are presented per mouse. GFP, green fluorescent protein; DAPI, 4′,6-diamidino-2-phenylindole; and RLU, relative luciferase units. Open circles, mice injected with saline (control group); closed circles/rectangles, mice injected with plasmids expressing Esm-1, luciferase, and transposase. R^2^ and p-values for simple linear regression (solid lines) are shown. Dotted lines indicated 95% confidence interval. * p < 0.05 vs. saline-injected mice at the indicated time points by one-way ANOVA. N= 3-5 mice per group.

### Mouse and Human Esm-1 decrease the degree of albuminuria in diabetic mice, independent of leukocyte infiltration

We performed hydrodynamic injection of mouse Esm-1 plasmid one week prior to the induction of diabetes (**Figure 4A**). Hydrodynamic injection significantly increases circulating plasma Esm-1 approximately ten-fold (1.4 ± 0.3 vs. 12.7 ± 1.5 ng/mL, p-value < 0.01, N= 10-11 mice per group)) (**Figure 4B**). Over-expression of mouse Esm-1 does not appreciably alter the incidence of hyperglycemia or alter body weight (**Figure 4D,E**). Esm-1 attenuates the induction of albuminuria in diabetic mice (220.2 ± 42.2 vs. 1193.1 ± 396.3 mg/g, p-value = 0.55, N= 10-15 mice per group) compared to saline-injected controls (261.0 ± 70.7 vs. 3165.9 ± 1308.5 mg/g, p-value < 0.01, N= 11-12 mice per group) (**Figure 4F**). In contrast to the effects on albuminuria, over-expression of mouse Esm-1 does not reduce either total glomerular leukocytes (**Figure 4G**) or tubulointerstitial leukocytes or specific leukocyte subsets in either kidney compartment (**Supplementary Figure 4**).

**Figure 4:**
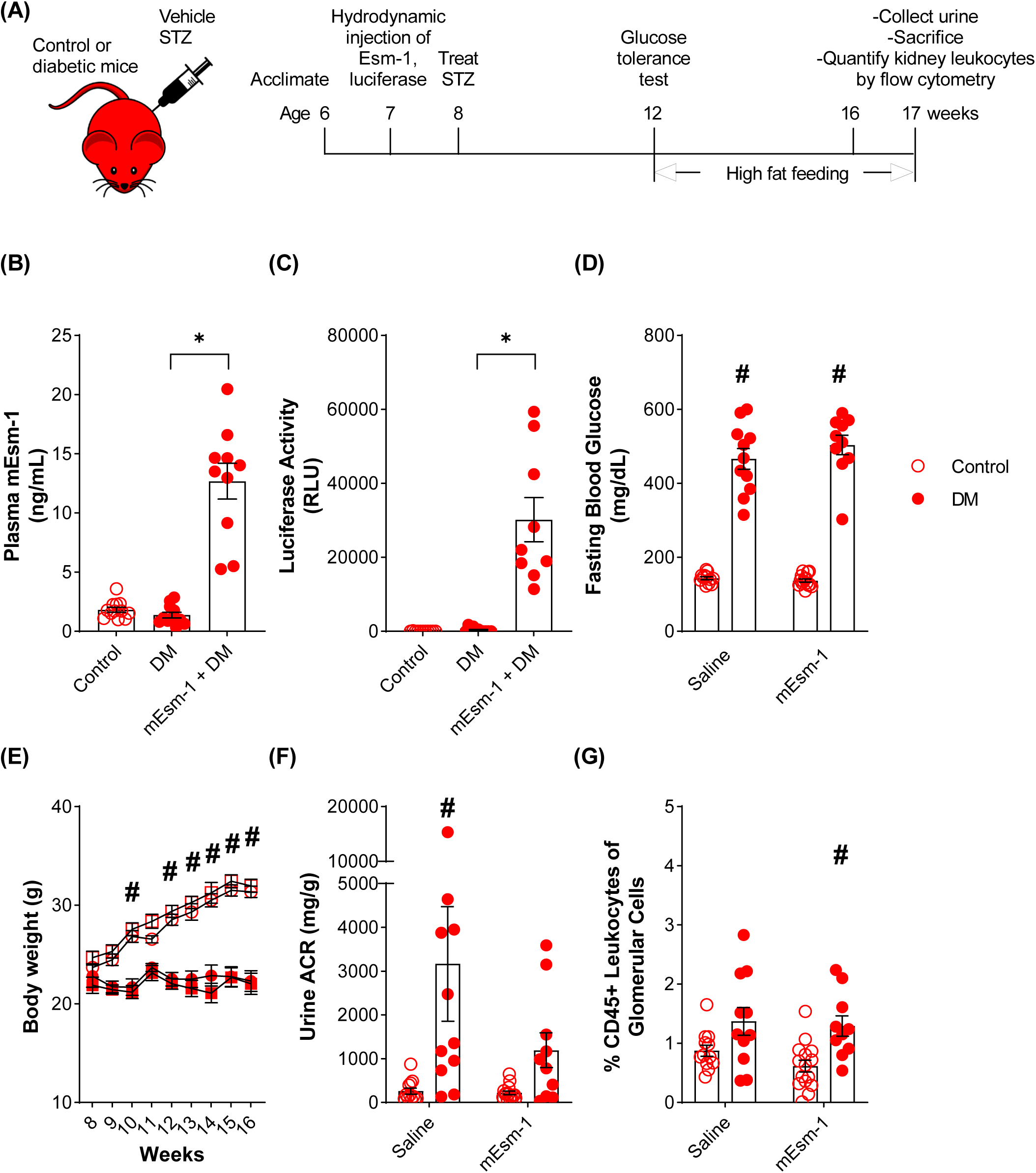
Systemic over-expression of mouse Esm-1 reduces diabetes-induced albuminuria in DKD-susceptible mice. (A) Timeline of experimental design. We induced expression of mouse Esm-1 (mEsm-1) by hydrodynamic injection and treated mice with vehicle vs. streptozotocin (STZ) and high fat feeding at indicated time points, followed by sacrifice at 17 weeks of age. We quantified (B) plasma Esm-1, (C) luciferase activity, (D) fasting blood glucose, (E) body weight, (F) urine albumin-to-creatinine ratio (ACR), and (G) glomerular leukocytes in control and diabetic mice. Open circles, control mice; closed circles, diabetic (DM) mice. In all panels except (E), results are presented per mouse and include mean ± SEM. In (E), values for mice with over-expression of mEsm-1 shown as squares and results are presented as mean ± SEM. RLU, relative luciferase units. * p-value < 0.05 compared across groups as indicated; # p-value < 0.05 vs. control mice, by one-way ANOVA (B-C) or two-way ANOVA (D-G). N= 9-15 mice per group.

We also performed hydrodynamic injection of human Esm-1 plasmid one week prior to the induction of diabetes (**Supplementary Figure 5A**). Hydrodynamic injection significantly increases circulating plasma human Esm-1 (0.03 ± 0.03 vs. 5.61 ± 1.41 ng/mL, p-value = 0.02, N= 8-10 mice per group) (**Supplementary Figure 5B**). Over-expression of human Esm-1 attenuates the induction of albuminuria in diabetic mice (103.0 ± 23.4 vs. 118.0 ± 57.3 mg/g, p-value > 0.99, N= 6-8 mice per group) compared to saline-injected controls (99.2 ± 41.5 vs. 690.0 ± 300.0 mg/g, p-value = 0.03, N= 11-12 mice per group) (**Supplementary Figure 5D**). In contrast to the effects on albuminuria, over-expression of human Esm-1 also does not reduce either total glomerular leukocytes (**Supplementary Figure 5E**) or tubulointerstitial leukocytes or specific leukocyte subsets in either kidney compartment (**Supplementary Figure 6**). Taken together, the glomerular responses to Esm-1 are consistent across species.

### Esm-1 deletion in DKD-resistant mice increases albuminuria, independent of leukocyte infiltration

As a complementary approach, we engineered and validated a constitutive knockout of Esm-1 on a DKD-resistant background (**Figure 5A,B**). Knockout mice have no detectable circulating plasma Esm-1 and with induction of diabetes, have similar fasting blood glucose and body weight compared to DKD-resistant wild-type littermate controls (**Figure 5D-F**). The degree of diabetes-induced albuminuria is modestly but significantly higher in Esm-1 knockout mice (50.6 ± 14.6 vs. 296.8 ± 65.8 mg/g, p-value < 0.01, N= 8-11 mice per group) compared with DKD-resistant wild-type littermate controls (38.1 ± 8.8 vs. 207.0 ± 100.5 mg/g, p-value = 0.06, N= 6-9 mice per group) (**Figure 5G**). In contrast to the effects on albuminuria, Esm-1 knockout mice do not exhibit diabetes-induced glomerular leukocyte infiltration (**Figure 5H**). In fact, glomerular leukocytes (macrophages) are modestly decreased in knockout mice in diabetics vs controls. Tubulointerstitial leukocytes or leukocyte subsets are not appreciably different in knockout mice or wild-type littermate controls (**Supplementary Figure 7**).

**Figure 5:**
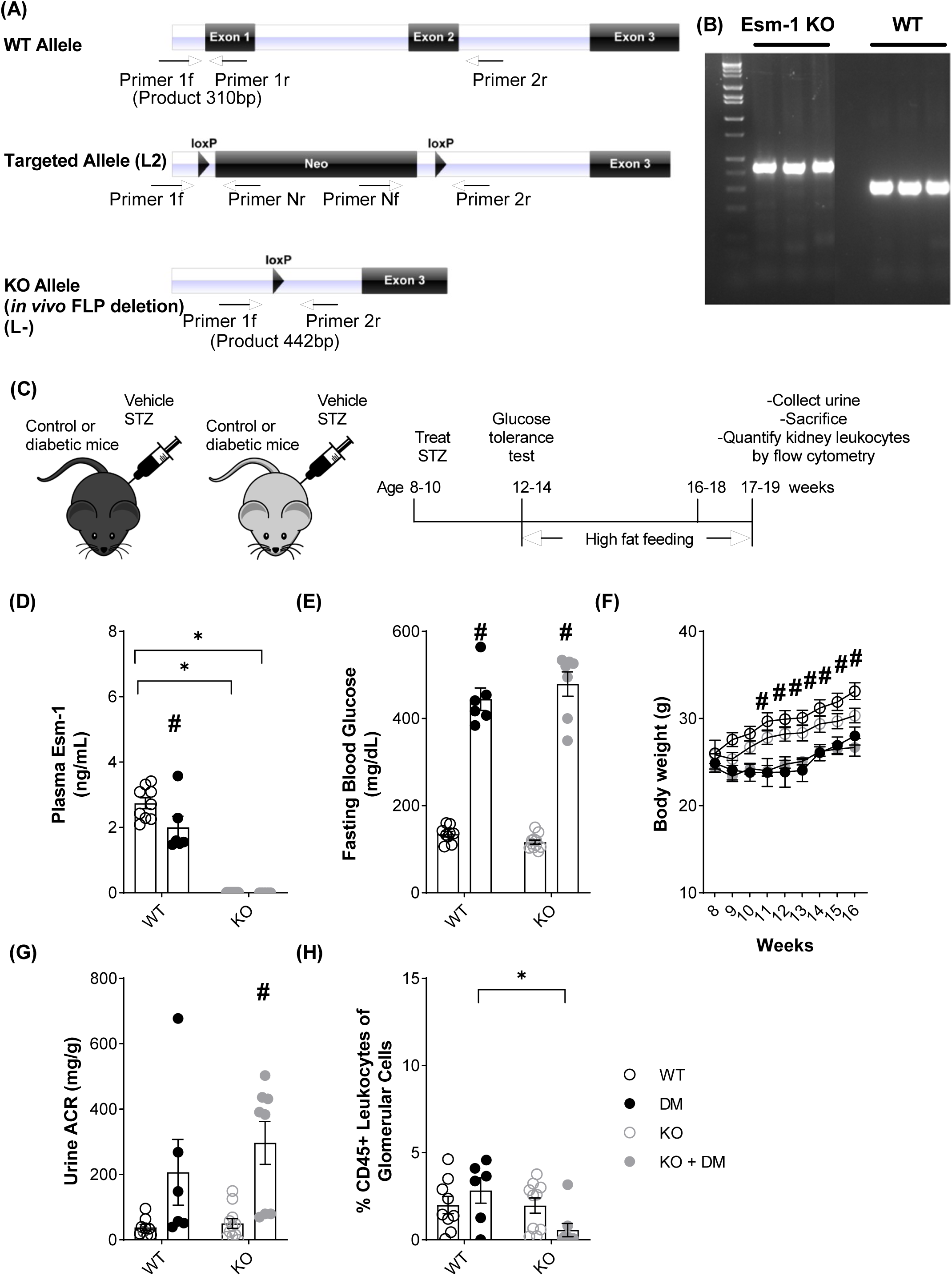
Constitutive deletion of Esm-1 induces albuminuria in diabetic DKD-resistant mice. (A) Schematic of DNA structure of wild-type (WT), targeted, and knockout (KO) alleles for generation of Esm-1 knockout mice. (B) DNA agarose gel showing results from genotyping of Esm-1 KO and WT littermate controls. (C) Timeline of experimental design. We treated wild-type and Esm-1 knockout mice with vehicle vs. streptozotocin (STZ) and high fat feeding, followed by sacrifice at 17-19 weeks of age. We quantified (D) plasma Esm-1, (E) fasting blood glucose, (F) body weight (G) urine albumin-to-creatinine ratio (ACR), and (H) glomerular leukocytes in control and diabetic, wild-type and Esm-1 knockout mice. Open circles, control mice; closed circles, diabetic (DM) mice. *Black*, wild-type (WT) littermate controls. *Grey*, Esm-1 knockout (KO) mice. In all panels except (D), results are presented per mouse and include mean ± SEM. In panel (D), results are presented as mean ± SEM. * p < 0.05 compared across groups as indicated; # p < 0.05 vs. control mice by two-way ANOVA. N= 6-11 mice per group.

### Esm-1 attenuates WT-1(+) podocyte injury

In histologic analysis, using WT-1 as marker of both parietal and visceral podocytes^25, 26^ (**Supplementary Figure 8**), diabetes significantly decreases the number of WT-1(+) podocytes (**Figure 6A**), and addition of mouse Esm-1 prevents this podocyte loss (**Figure 6B**). Using p57 as a marker primarily of visceral podocytes, diabetes significantly, but only modestly, increases p57(+) podocyte nuclear area (adjusted p-value < 0.001) (**Supplementary Figure 9C**) and glomerular podocyte coverage (adjusted p-value < 0.001) (**Supplementary Figure 9D**), markers of podocyte hypertrophy / injury^25, 27^. Over-expression of mouse Esm-1 significantly reduces glomerular p57(+) podocyte coverage compared to diabetes alone (p-value < 0.001) (**Supplementary Figure 9E**) (**Supplementary Table 1**). In contrast to WT-1, we find that p57(+) podocytes do not decrease at this stage of diabetes (**Supplementary Figure 9F**,**G**). Diabetes increases fractional glomerular area of fibronectin and collagen type IV, sensitive markers of mesangial matrix expansion^28^; however, over-expression of Esm-1 does not significantly reduce expression of these matrix proteins at the time of sacrifice compared to diabetes alone (**Supplementary Figure 10**). Taken together, over-expression of Esm-1 attenuates albuminuria in mice with DKD, possibly through preservation of podocyte health.

**Figure 6:**
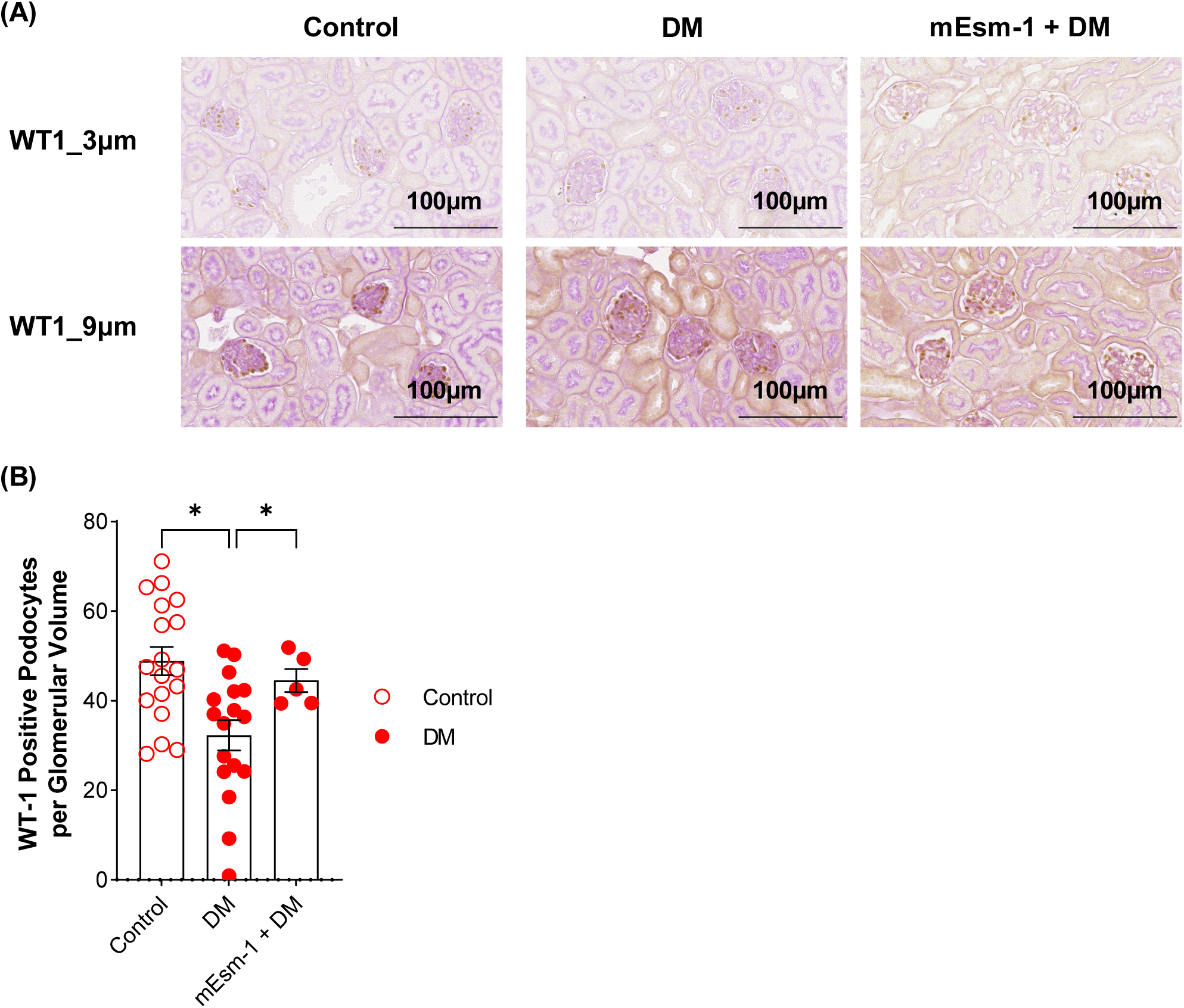
Systemic over-expression of mouse Esm-1 restores podocyte loss induced by diabetes. (A) Representative images of mouse kidney sections stained for WT-1 expression. Scale bars (*black*) as indicated. (B) WT-1(+)podocytes per glomerulus on larger cohort mice. N=5-18 mice per group. Open circles, control mice; closed circles, diabetic (DM) mice. * p < 0.05 compared across groups as indicated by one-way ANOVA.

### Esm-1 attenuates interferon signaling in diabetic kidney disease

To understand the mechanism of how Esm-1 significantly prevents diabetes-induced kidney injury, we explored transcriptomic responses in glomeruli from a representative subset of control, diabetic, and diabetic mice treated with Esm-1 (**Supplementary Figure 11**). Using bulk RNAseq, diabetic mice demonstrate several pathways that are up- and down-regulated (**Figure 7A**). Strikingly, of the top 11 significantly up-regulated pathways, seven of these relate to interferon signaling. When we compare control vs. diabetic mice with over-expression of mouse Esm-1, the majority of these pathways are not significantly up-regulated. When we analyze by interferon-related genes, 85 are differentially expressed in diabetic vs. control mice compared with only 19 between diabetic with over-express Esm-1 vs. control mice (**Supplementary Figure 12**). In an orthogonal analysis of the most up- or down-regulated genes between diabetes with over-expression of Esm-1 vs. diabetes alone, we detect 41 significantly and differentially expressed genes (**Figure 7B**). From this list, we note several solute carriers, but also honed in on genes with known association with kidney disease, *Ackr2*^29^ and *Cxcl11*^30^. *Ackr2* and *Cxcl11* are significantly down-regulated with over-expression of Esm-1 (**Figure 7C,D**), and we validate these findings by qPCR in a larger cohort of mice (**Figure 7E,F**).

**Figure 7:**
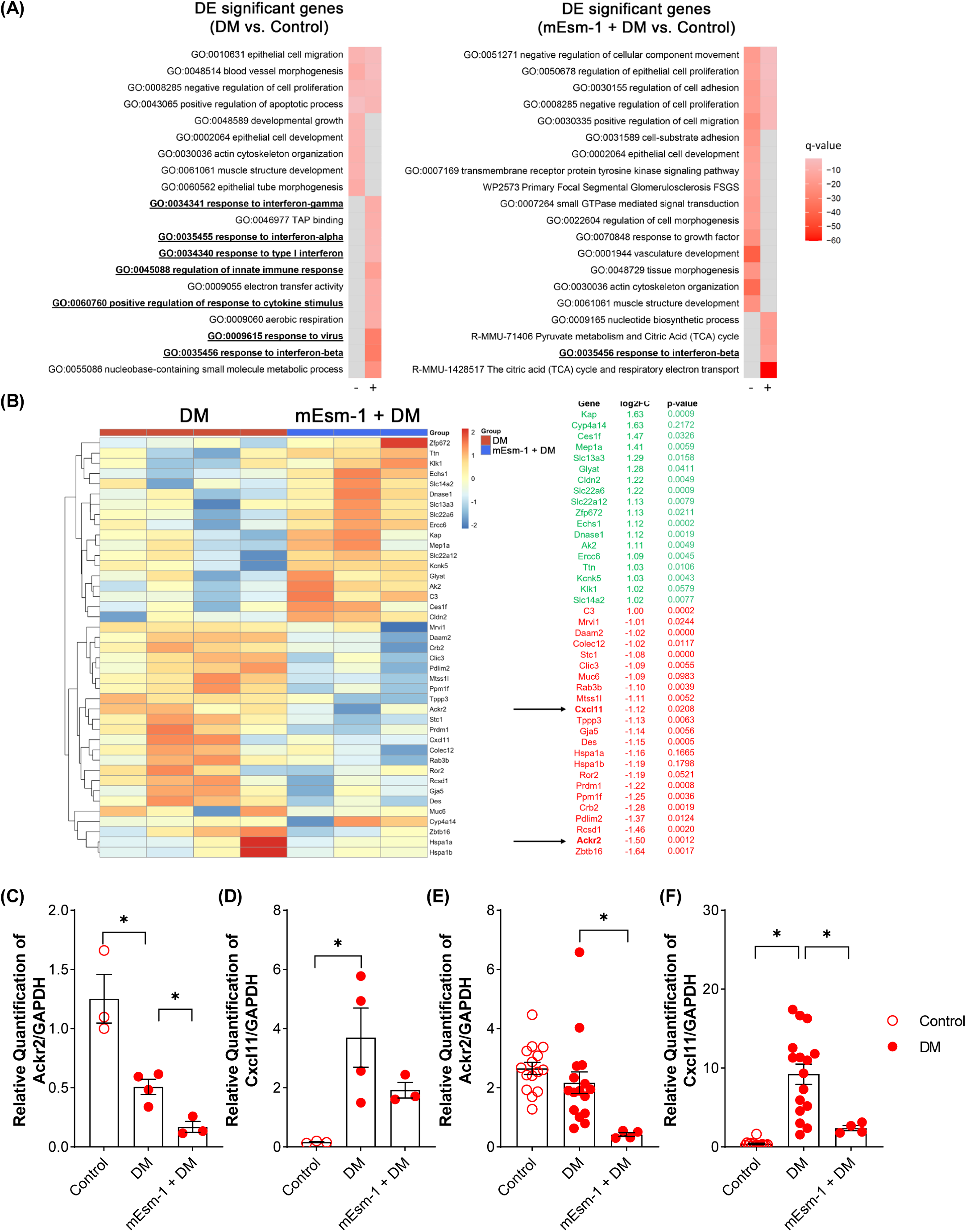
Esm-1 attenuates chemokine/chemokine receptor expression in glomeruli. (A) Heatmaps of enrichment of ontology clusters, showing the hypergeometric q-values of gene ontologies falling into each term for the top 20 most significantly enriched ontology clusters in each gene list. We performed a hierarchical clustering of significant ontology terms into a tree based on Kappa-statistical similarities among their gene memberships. We selected the term with the best q-value within each differentially-expressed (DE) cluster as its representative term and display it in the heatmaps. Grey heatmap cells represent the non-significantly enriched ontology clusters in a given gene list. Light red is lowest q-value, dark red is highest q-value. +: Pathways up-regulated with diabetes or diabetes + mouse Esm-1, relatively to control; -: Pathways down-regulated with diabetes or diabetes + mouse Esm-1, relatively to control. (B) *Left panel*, Heatmap with hierarchical clustering of glomerular genes significantly differentially regulated in diabetic (DM) mice vs. diabetic mice with over-expression of mouse Esm1 (mEsm-1 + DM); Colors in cells from individual mice represent z-scores relative to mean of DM group, scale on right; *Right panel*, Genes up-(*green*) or down-regulated (*red*) in glomeruli from diabetes (DM) vs. diabetes + mouse Esm-1 (mEsm-1 + DM) with log_2_ fold change (FC), p-values, and q-values. Arrows highlight chemokine/chemokine receptor genes, *Ackr2* and *Cxcl11*. (C-D) Validation by qPCR of *Ackr2* (D) and *Cxcl11* (D) expression relative to GAPDH in samples used for glomerular RNAseq. * p-value < 0.05 as indicated. N= 3-4 mice per group. (E-F) Validation by qPCR of *Ackr2* (E) and *Cxcl11* (F) expression relative to GAPDH in all samples from mouse over-expression experiments, phenotyped for clinical and histologic characteristics. * p-value < 0.05 as indicated. N= 9-15 mice per group.

## Discussion

Esm-1 transcription is stimulated by pro-inflammatory cytokines^18^, and progressive DKD is characterized by inflammation^15, 31-35^. Although others have reported correlations of Esm-1 with kidney diseases^23, 36-38^, cardiovascular diseases^23, 39-42^, and cancer^43-47^, data linking Esm-1 to DKD are mixed. For example, Ekiz-Bilir et al. showed a direct correlation between serum Esm-1 and the degree of albuminuria in patients with type 2 diabetes mellitus^48^, while Cikrikcioglu et al. reported the opposite^49^. Some studies have even shown a deleterious effect of Esm-1, albeit in vitro^50^. Glomerular Esm-1 is relatively deficient in patients and DKD-susceptible mice^13^, implicating Esm-1 as a possible protective gene. Using cross-sectional data, we do not observe a lower circulating level of Esm-1 in patients with diabetes and albuminuria (**Supplementary Figure 1**), However, in comparing longitudinal outcomes in patients with minimal initial kidney injury, circulating Esm-1 was lower in those patients that developed progressive stages of albuminuria (**Figure 1**). These data are limited by small sample size, potential confounding, and short duration of follow-up. In future work we will validate these findings in larger, longitudinal cohorts and by major adverse kidney events such as changes in eGFR or onset of renal replacement therapy. In DKD-susceptible mice, we found that, Esm-1 > 3 ng/mL is associated with protection from albuminuria or glomerular leukocyte infiltration (**Figures 2,3**). In mice, we did not have data on plasma Esm-1 concentrations prior to the onset of kidney disease. Taken together, these data suggest at least two potential interpretations: (1) Esm-1 protects from worsening DKD; or (2) Esm-1 is a marker but not a mediator of DKD. To distinguish between these possibilities and to determine causality in vivo, we employed complementary approaches to over-express Esm-1 in DKD-susceptible mice and delete Esm-1 from DKD-resistant mice.

We find that Esm-1 has renoprotective effects. DKD-susceptible mice have diabetes-induced albuminuria, glomerular leukocyte infiltration, podocyte injury, and increased production of extracellular matrix proteins. We utilize Esm-1 over-expression to assess its contribution to these sequelae of kidney injury. Our data suggest that mouse and human Esm-1 reduces albuminuria independent of glomerular leukocyte infiltration (**Figure 4, Supplementary Figure 5**). To complement gain-of-function experiments on a DKD-susceptible background, we utilized a transgenic mouse model for loss of function of Esm-1 on a DKD-resistant background. This result is congruent with results from over-expression experiments; however, the effect of Esm-1 deletion on DKD-resistant background is more modest (**Figure 5**). While one may anticipate that Esm-1 deletion would not reduce glomerular leukocyte infiltration, in constitutive knockout mice, we must account for the role of Esm-1 in vascular development^20^. Indeed, Rocha *et al*. showed that constitutive Esm-1 knockout mice had a higher transcellular resistance, and in an acute model of leukocyte infiltration, were also protected from inflammation. Thus, the present study is the first to demonstrate that Esm-1 is sufficient for renoprotection in DKD (**Figure 8**).

**Figure 8:**
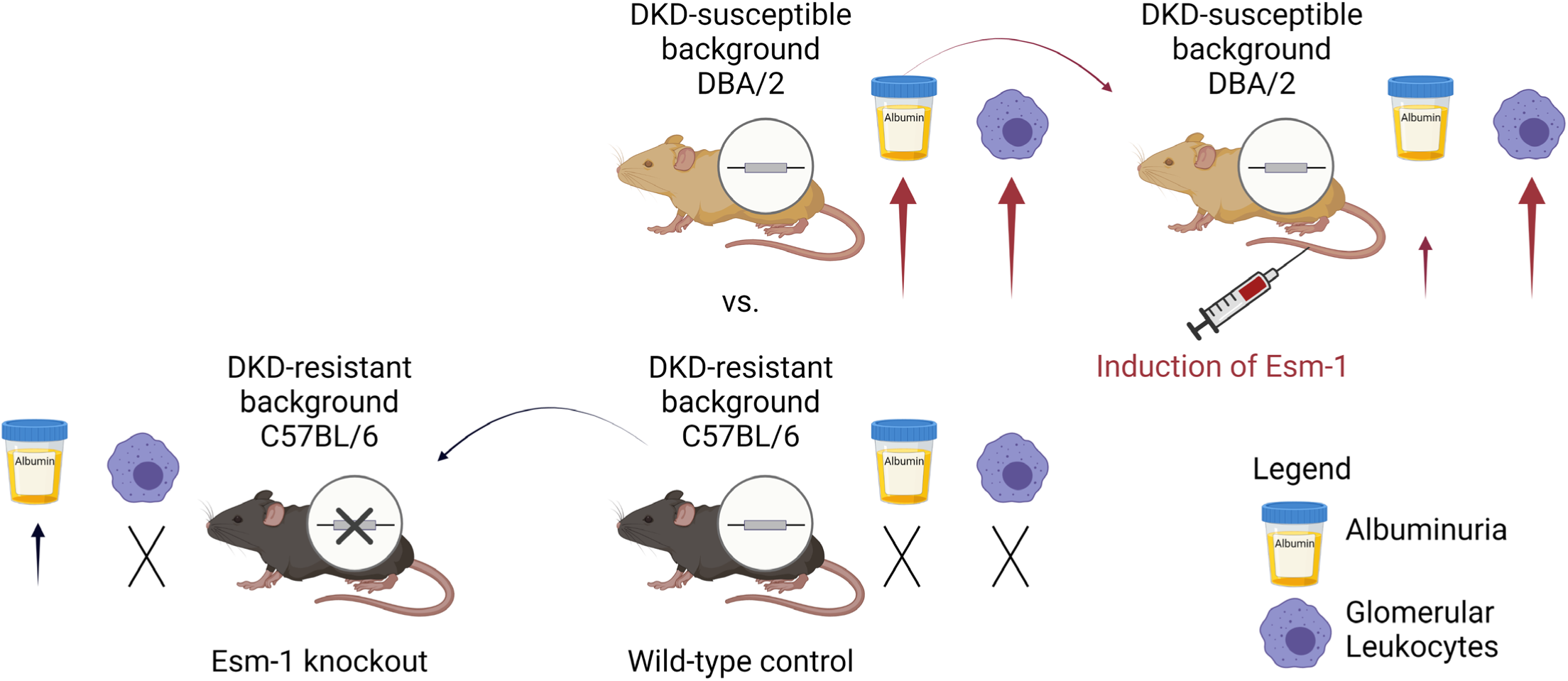
Overview of results with gain- or loss-of-function of Esm-1 in diabetic mice. In DKD-susceptible mice (DBA/2), diabetes induces albuminuria and glomerular leukocyte infiltration in contrast to DKD-resistant mice (C57BL/6), with minimal albuminuria or infiltration. Induction of mouse or human Esm-1 in DKD-susceptible mice reduces albuminuria, independent of leukocyte infiltration. In contrast, Esm-1 deletion from DKD-resistant mice modestly increases albuminuria but not leukocyte infiltration. Arrows (↑) or ‘X’ symbols indicate degree or absence of kidney injury, respectively, in diabetic mice relative to controls. DKD, diabetic kidney disease.

As evidenced by in vitro and in vivo studies, the anti-inflammatory mechanisms of Esm-1 are well-described in other systems but have not been shown in glomeruli or in chronic conditions such as diabetes. Esm-1 treatment both in vitro and in vivo reduces leukocyte infiltration acutely. Esm-1 binds to LFA-1 on Jurkat cells, and blocks LFA-1-ICAM-1 interaction^19^. We have previously shown that Esm-1 inhibits human leukocyte transmigration, but not adhesion in a microfluidic flow chamber assay ex vivo^13^. Intraperitoneal injection of human Esm-1 in Balb/c mice one hour after lipopolysaccharide administration mildly but significantly decreased lung neutrophil infiltration *in vivo*^17^. However, our results show that Esm-1 inversely correlates with but does not inhibit leukocyte infiltration in mice several weeks after the onset of diabetes. Possible explanations for this observation, include the following: (1) constitutive Esm-1 may determine leukocyte infiltration, and we over-expressed Esm-1 in adulthood; (2) Esm-1 may reduce distinct subtypes of macrophages^51^ and further leukocyte phenotyping is needed; (3) Esm-1 only alters the amount of resident leukocytes (e.g. macrophages); and/or (4) compensatory mechanisms overcome an acute effect of Esm-1, and long-term effects of Esm-1 are on structural features of DKD downstream of infiltrating leukocytes.

Using WT-1, p57, and podometrics, we show that Esm-1 significantly reduces the total number of WT-1(+) podocytes in a mouse model of diabetes, consistent with prior literature^52^. Esm-1 also modestly reduces p57(+) podocyte nuclear area, a marker of podocyte hypertrophy^27^. Our diabetic mice also show modest increases in extracellular matrix proteins-fibronectin and collagen type IV. Collectively, these data suggest that we assayed for the effects of Esm-1 at an early stage of DKD^53^. Further studies are needed to determine whether sustained over-expression of Esm-1 may reduce additional sequelae of DKD.

Endogenous Esm-1 is primarily localized to glomeruli, and DKD-susceptible mice have a deficiency of Esm-1 production specifically in glomeruli^13^. While specific over-expression from glomerular endothelial cells was not technically feasible, in the present study, we used hydrodynamic injection to over-express Esm-1 and increase circulating Esm-1. Esm-1 is a locally derived inhibitor of inflammation and manipulating the level of this inhibitor at the local level is an attractive candidate approach to attenuate DKD without the disadvantages of systemic immunosuppression. Our observation of different kidney phenotypes and transcriptomic changes in glomeruli, implies direct effects of circulating Esm-1 on kidney, and motivates future studies to over-express Esm-1 specifically in glomerular endothelial cells and across multiple mouse models of DKD and possibly other podocytopathies.

A glomerular signature of interferon-inducible genes that we observe in diabetic vs. control mice is pathophysiologically relevant for DKD. Several cytokine cascades in the pathogenesis of DN including TNF-α, IL-6, and IL-18^54, 55^ implicate interferon signaling. IL-18 is a predictor of albuminuria in T2DM and DN^56, 57^ and higher IL-18 levels are associated with more severe DN^58-60^. The Nrlp3 inflammasome, that increases mature IL-18 aggravates DN^61^. IL-18 is a prominent inducer of interferon-gamma via NF-κB^62, 63^, and NF-κB inhibition attenuates diabetic kidney injury^64^. Interferon-gamma is up-regulated in patients with diabetes and kidney disease compared with diabetes alone^65^, and interferon-gamma also up-regulates the inflammasome^66^. Toll-like receptors are activated in human kidney and mediate the pathogenesis in rodent models of DN^67-69^. When stimulated, these toll-like receptors stimulate gene transcription via NF-κB such as the NF-κB target gene, type 1 interferon-beta^70^. Transcription of interferon-inducible genes are then synergistically stimulated by NF-κB and interferon signaling. Notably, genes up-regulated in progressive human DKD share a well-recognized NF-κB- and interferon-response element (NFKB_IRFF_01)^71, 72^. Therefore, one role for Esm-1 to mitigate DKD may be through its ability to attenuate the transcription of interferon-inducible genes. Interestingly, Esm-1 transcription is stimulated by several cytokines but is inhibited by interferon-gamma^18^. Based on our data, one consequence of interferon-gamma-mediated reduction of Esm-1 transcription may be more effective propagation of interferon signaling.

Another aspect of inflammation in DKD is up-regulation of chemokines and adhesion molecules leading to leukocyte recruitment^54, 69^. However, direct effects on glomerular cell types (e.g. podocyte apoptosis, mesangial matrix accumulation, and endothelial cell permeability) are also well-known characteristics of an inflammatory phenotype^73, 74^. For example, toll-like receptors and interferons can mediate podocyte apoptosis^75-77^. Moreover, the chemokine receptor CCR2, on macrophages, mediates leukocyte infiltration, but the same gene activated specifically on podocytes, mediates podocytes apoptosis, independent of macrophage infiltration^74^. Chronic over-expression of systemic Esm-1 mediates albuminuria and structural features of DKD but not inflammatory cell infiltration. Thus, the mechanisms of Esm-1 provide an example of separate but related pathways in DKD. Albuminuria and glomerular leukocyte infiltration did not correlate in our mouse model (**Supplementary Figure 13**). In prior human studies, leukocyte infiltration and albuminuria each correlate directly with tubulointerstitial fibrosis, but do not correlate directly with each other^31, 32^, supporting the possibility that these important sequelae of kidney injury, at least in early DKD, may be, in part, due to separate mechanisms.

We validated selected target genes as potential mediators of the effects of Esm-1 on glomeruli. Two genes, *Ackr2* and *Cxcl11*, belong to a family of chemokines and chemokine receptors implicated in kidney disease and interferon signaling^78-82^. *Ackr2* can internalize chemokines for degradation which reduces chemokine availability, inhibits leukocyte infiltration, and promotes immune resolution and adaptive immunity. However, the functions of *Ackr2* may be context-dependent, and may be pathologic^29, 83, 84^ or protective^29^ in different disease models^85^. Genetic deletion of *Ackr2* attenuates albuminuria, leukocyte infiltration, and interferon-related pathways (e.g. NF-κB and pattern recognition receptors) in diabetic mice^29^, suggesting loss of function of *Ackr2* is renoprotective in susceptible mice models of DKD^29^. We similarly show that over-expression of Esm-1 down-regulates glomerular *Ackr2* in diabetic DKD-susceptible mice, with attenuated albuminuria and attenuated interferon signaling. *Cxcl11* and its receptor, CXCR3, mediate chemotaxis and proliferation of immune cells in interferon-induced inflammation^81, 82, 86^. In patients with chronic kidney disease, urine CXCL11 is higher in later stages of DKD and correlates with progression of disease^30^. Additional ligands for CXCR3, are up-regulated in sera from patients with DKD vs diabetes alone^59^. In a mouse model of acute glomerular inflammation, systemic genetic deletion of CXCR3 attenuates glomerulosclerosis and albuminuria^87^. Notably, *Cxcl11* is not functional in DKD-resistant mice (C57BL/6)^88^ due to insertion of two nucleotides and a resultant premature stop codon and nonsense-mediated decay. Different mouse strains susceptible to DKD including DBA/2 and BALB/c^8, 9^, produce a functional copy of *Cxcl11*. A non-functional allele of *Cxcl11* in C57BL/6 mice may be an explanation for why Esm-1 deletion may only have a modest permissive effect on albuminuria on this background strain, as deletion could not reciprocally increase *Cxcl11*. In future studies to assess the protective role of Esm-1, we will characterize the contribution of these chemokine-related genes and other differentially regulated genes in interferon-dependent and interferon-independent pathways.

Integration of our histologic and transcriptomic data, suggests that Esm-1 may more potently attenuate the parietal vs. visceral podocyte apoptosis of diabetes^89, 90^. We show that Esm-1 attenuates multiple interferon-stimulated glomerular genes, including *Cxcl11*, but not genes of the interferon-beta pathway. Interferon-alpha, compared to interferon-beta, markedly stimulates kidney *Cxcl11*^77^. Interferon-alpha, in contrast to -beta, also induces apoptosis in parietal vs. visceral podocytes in vitro and in vivo^77^. In histologic studies, differences in the protective effect of Esm-1 on WT-1(+) vs. p57(+) podocytes, may be driven by effects on parietal podocytes, as WT-1 expression is more pronounced in these cells compared to p57 (**Figure 6, Supplementary Figures 8-9**). Thus, Esm-1-mediated attenuation of interferon-alpha and *Cxcl11* may be one mechanism to preserve parietal podocyte integrity and to reduce albuminuria. In future studies, we will measure the contribution of Esm-1 in the presence or absence of *Cxcl11*.

In summary, Esm-1 predicts protection from DKD in humans and inversely correlates with DKD in mice. We determine that Esm-1 rescues DKD-susceptible mice from podocyte injury and albuminuria, and Esm-1 deletion is sufficient to partially reverse the protective background of DKD-resistant mice. Esm-1 attenuates interferon-stimulated pathways and down-regulates chemokine signaling, e.g., glomerular *Acrk2* and *Cxcl11*, which are part of a potential mechanism to attenuate DKD. Taken together, Esm-1 appears to be renoprotective in mice and humans, and up-regulation of Esm-1 should be explored as a possible therapeutic strategy in DKD.

## Materials and Methods

### Biorepository

We collected serum samples from patients in two separate cohorts: (1) Patients who self-identified with diabetes mellitus at the Palo Alto Medical Foundation from 2011-2012 (N= 250); and (2) patients who were consented for the Stanford Precision Biobank regardless of status of diabetes mellitus or kidney disease (N=2579). From these cohorts, we extracted data from the electronic health record and maintained a de-identified, secure database. We verified patients with type 2 diabetes using physician recorded diagnosis (ICD-9 codes 250.X0, 250.X2), abnormal laboratory values according to American Diabetes Association guidelines [two of any hemoglobin A1c ≥ 6.5%, fasting blood glucose ≥ 126 mg/dL, random blood glucose ≥ 200 mg/dL, or oral glucose tolerance test ≥ 200 mg/dL]^91^ or use of any anti-diabetic medications. Among patients with at least one recorded urine albumin or protein and one serum creatinine measurement (N= 143 samples), after verification of diabetes mellitus, we classified these patients with: (a) normoalbuminuria, < 30, (b) microalbuminuria, 30-300; or (c) macroalbuminuria, >= 300 mg urine albumin per gram of creatinine. We retained patients in the cohort with a stable classification of albuminuria within the months immediately preceding and following the biospecimen collection date. For (1), a subset of patients with normoalbuminuria were matched to patients with micro- or macroalbuminuria by age, sex, and reported race/ethnicity. In total, we studied N=99 patients with sufficient and stable baseline characteristics. Among patients with normo- or microalbuminuria with a baseline eGFR ≥ 30 mL/min/1.73m^2^ using the CKD Epidemiology Collaboration equation^92^, we classified patients as progressors if longitudinal data showed a persistent transition from normo- to micro- or macroalbuminuria or micro- to macroalbuminuria. The time between baseline urine measurement and serum sample collection was (6 (13.5) months, median (IQR)) and follow-up data was available for 1.75 (3.0) years, median (IQR). Years of available follow-up data did not significantly differ between groups. We processed samples by centrifugation and stored aliquots at −80°C.

### Esm-1 ELISA

We quantified human serum Esm-1 by ELISA (Lunginnov, Lille, France) and mouse plasma Esm-1 by ELISA (Aviscera Biosciences, Santa Clara, CA), following the manufacturer’s instructions.

### Induction of diabetes

We injected 8-week old male DKD-susceptible (DBA/2J) or DKD-resistant (C57BL/6J) mice (Jackson Laboratories, Bay Harbor, ME) intraperitoneally with 40 mg/kg body weight of streptozotocin (STZ, Sigma-Aldrich, St. Louis, MO) vs. vehicle (sodium citrate [pH 4.5]) daily for five consecutive days. We then supplemented mice with 10% sucrose water to prevent hypoglycemia. Four weeks after the injection, we fed diabetic mice with a high fat diet (Research Diets, New Brunswick, NJ) for nine weeks until sacrifice^12, 93^. We measured body weight once per week. We quantified fasting blood glucose with Bayer Contour glucometer at four weeks post STZ injection to verify hyperglycemia. We collected urine samples at nine weeks post STZ injection and sacrificed mice at 10 weeks post STZ injection by cardiac perfusion. We prepared plasma samples from retro-orbital blood and stored them at −80°C. We then prepared glomerular and tubulointerstitial single cells for flow cytometry analysis.

### Quantification of Albuminuria

We collected urine from mice in singly housed in metabolic cages (Tecniplast, Italy). We quantified urine albumin by ELISA and creatinine by a Creatinine Companion kit (Exocell, Philadelphia, PA), following the manufacturer’s instructions. For quantification of albumin and creatinine, we diluted urine samples at 1:10 - 1:15 for vehicle-treated mice and at 1:100 - 1:300 for polyuric, diabetic mice.

### Single cell preparation of glomerular and tubulointerstitial tissue

We anesthetized mice with isoflurane and sacrificed them after cardiac perfusion. We isolated mice kidneys and dissociated tissue by forceps, and then by gentleMACS tissue homogenizer with the program h_cord 01.01 (Miltenyi Biotec, San Diego, CA) in 5 mL Hepes Ringer buffer (118 mM NaCl, 17 mM H-Hepes, 16 mM Na-Hepes, 14 mM Glucose, 3.2 mM KCl, 2.5 mM CaCl_2_·2H_2_O, 1.8 mM MgSO_4_·7H_2_O, 1.8 mM KH_2_PO4). After centrifugation (200 g for 10 minutes), we resuspended the kidney tissue in Hepes Ringer buffer containing 0.2% collagenase I (Worthington Biochemical, Lakewood, NJ) and 0.2% hyaluronidase (Sigma-Aldrich), 90 mL/g kidney weight, and rotated at 37°C for two minutes. We washed the digested tissue multiple times with Hepes Ringer and centrifuged at 30 g for two minutes, until the supernatant became clear. We resuspended the tissue in 13 mL Hepes Ringer buffer and passed tissue through a 40 μm cell strainer (Corning, Corning, NY) twice and a 100 μm strainer (Thermo Fisher Scientific, Waltham, MA) once to isolate glomeruli. The flow-through contained tubulointerstitial single cells. We centrifuged both glomerular and tubulointerstitial fractions at 4000 rpm for 10 min. We resuspended the tubulointerstitial single cells with 1 mL 4% neutral buffered formalin (Thermo Fisher Scientific) and fix for 20 minutes at room temperature, then stored in PEB buffer (1x PBS, 2 mM EDTA, 0.5% BSA) at 4°C. We resuspended the glomerular fraction in 1mL HBSS buffer with Ca^2+^ and Mg^2+^ (Hyclone, Logan, UT) containing 0.1-0.2 mg/mL Liberase^™^Research Grade (Roche, Mannheim, Germany), and incubated glomeruli in 24-well tissue culture plates at 37°C for 24 hours. The next day, we broke the glomeruli by pipetting up and down for 10 times and centrifuged the glomerular single cells at 8000 rpm for 10 minutes, fixed cells with 4% neutral buffered formalin (Thermo Fisher Scientific) for 20 minutes at room temperature, and then stored cells in PEB buffer at 4°C.

### Flow Cytometry

We permeabilized formalin-fixed glomerular and tubulointerstitial single cells with 0.25% Tween-20 in PEB buffer at room temperature for 30 minutes and blocked cells with Fc Block (clone 2.4G2, BD Biosciences, San Jose, CA) diluted in 0.25% Tween-20 on ice for 10 minutes. We stained cells with fluorescent conjugated antibodies (**Supplementary Table 2**) on ice for 30 minutes. After washing, we resuspended cells in 200 μL PEB buffer for flow cytometry analysis on a Scanford Flow Cytometer in the Stanford Shared FACS Facility. We used Flowjo 10 software (Becton Dickinson, Ashland, OR) to analyze data. For CD45(+) cells, CD68(+)Ly6G(-) cells were classified as macrophages, CD68(-)Ly6G(+) as neutrophils, and CD68(-)Ly6G(-) cells as others (likely lymphocytes).

### Hydrodynamic Injection to Over-express Circulating Esm-1

Dr. Justin Annes (Stanford University) kindly provided pT3-hEsm-1-V5 and pT3-GLuc (*Gaussia* Luciferase). The Emory University Integrated Genomics core facility (Atlanta, GA) subcloned mouse and human Esm-1 cDNA from pCMV6 (Origene, Rockville, MD) into pT3. We prepared plasmids using EndoFree Plasmid Maxi Kit (Qiagen, Germantown, MD). One week before the induction of diabetes, we performed hydrodynamic injection to over-express Esm-1. We diluted three plasmids: pT3-mEsm-1-V5 (25 μg), pT3-GLuc (25 μg), and pCMV(CAT)T7-SB100 Sleeping Beauty transposase (Addgene, Cambridge, MA^94^, 2.5 μg) in endotoxin-free saline (Teknova, Hollister, CA). We injected the plasmid solution through the mouse tail vein at 0.1 mL per gram body weight with a 27G needle in 6-8 seconds. Saline injection serves as the negative control. At the time of sacrifice, we collected mice collected retro-orbital blood, prepared plasma samples, and quantified Esm-1 and luciferase activity. To validate the uptake of plasmids by hepatocytes, we performed hydrodynamic injection to express GFP. We sacrificed these mice and verified green fluorescence protein expression using a Leica DM5000 (South San Francisco, CA).

### Gaussia Luciferase Assay

We quantified plasma luciferase activity by a BioLux *Gaussia* Luciferase Assay Kit (NEB, Ipswich, MA) from 10 μL plasma acquired by retro-orbital bleeding, following the manufacturer’s instructions.

### Generation of Esm-1 knockout mice

We inserted loxP recognition sites to flank exons 1 and 2 of the mouse Esm1 gene (Ensembl ID ENSMUSG00000042379) (**Figure 5**). In collaboration with the Institut Clinique de la Souris (Strasbourg, France), we transfected a targeting cDNA fragment containing a neomycin resistance cassette surrounded by loxP sites into mouse embryonic stem cells by electroporation, allowing replacement of exon 1 and 2 of Esm-1 by the neomycin resistance cassette through homologous recombination. After transfection, we selected embryonic stem cells for recombination of Esm-1 with G418-containing medium, validated these clones by southern blot, and injected clones into blastocysts and then into pseudopregnant female mice. We then cross bred the offspring to generate mice that lack Esm-1 exon 1 and 2 on homologous chromosomes. We then cross bred these mice with Cre deleter mice to remove the neomycin resistant cassette. For genotyping, we extracted genomic DNA from mouse tails and performed PCR with REDExtract-N-Amp^™^ Tissue PCR Kit following the manufacturer’s instructions (Sigma-Aldrich) and analyzed samples on agarose gel. We utilized 1f and 1r primer pairs to generate a 310bp fragment for the wild-type allele and 1f and 2r primer pairs to generate a 442bp fragment, indicative of the knockout allele. Primer sequences are in **Supplementary Table 3**.

### Podocyte Staining

We stained three and nine μm paraffin kidney sections with anti-WT1 antibody (ab15249, Abcam, Cambridge, MA, USA) according to published protocols^95^. Briefly, we performed antigen retrieval on rehydrated sections in sodium citrate buffer pH 6.0 at 95°C for 1h. After blocking with 20% goat serum, we incubated sections with 1:400 diluted anti-WT1 antibody at 4°C overnight. We then blocked sections with a Biotin/Avidin blocking kit, incubated with biotinylated goat anti-rabbit secondary antibody, and developed with ABC and DAB kits (Vector Lab, Burlingame, CA, USA). We further stained sections with periodic acid and Schiff reagents (Sigma-Aldrich), without hematoxylin counterstain. After dehydration, we mounted sections with EUKITT mounting medium (Thermo Fisher Scientific). We captured images of stained sections with Nanozoomer digital scanner (Stanford Canary Center Imaging Core Facility). We then exported images containing 50 glomeruli per mouse and analyzed with machine learning tools developed with the Cord online platform (https://app.cord.tech/login). We used approximately 40 images to train models to initiate the recognition of glomerular tuft area. We employed a confidence of 0.4 to recognize podocytes and 0.6 to recognize the glomerular tuft area. We then quantified podocyte per glomerular volume according to the AMDCC protocol.

### RNA sequencing

We extracted kidney glomeruli from control, diabetic, and diabetic mice with over-expression of mouse Esm-1. We isolated glomerular RNA by the Zymoresearch RNA preparation micro kit (Zymoresearch, Irvine, CA). We removed genomic DNA by RNase-free DNase I (Fermentas, Waltham, MA). We then reversed transcribed RNA into cDNA by using Improm II reverse transcriptase (Promega, Madison, WI) and random hexamer primers (Genelink, Elmsford, NY). We submitted cDNA samples to CD Genomics (New York, NY) for DNA quality control, library preparation, and sequencing (Illumina Novaseq PE150 sequencing, 20M read pairs). We then trimmed raw paired-end FASTQ files using Trimmomatic v0.36. We performed RNA-Seq alignment to the reference mm9 using STAR v2.5.3a and default conditions. We filtered gene counts for lowly expressed genes and performed differential expression analyses using the DESeq2 v1.28.1 R package installed in R v4.0.2, with default settings. We submitted the data to GEOBANK with accession number: GSE175449.

### RT-qPCR

We prepared cDNA as described above. We then used Fast Syber Green Master Mix (Applied Biosystems, Foster City, CA) to quantify cDNA samples (1:5 – 1:25 diluted) with the primers listed in **Supplementary Table 4** at a melting temperature of 60°C. We used the StepOne PCR machine (Applied Biosystems, Foster City, CA) provided by the Stanford Protein and Nucleic Acid (PAN) Facility.

### Statistics

For statistical comparisons with normal data distribution, we used unpaired Student’s t-test as appropriate for two columns, parametric one-way ANOVA as appropriate for more than two columns, and two-way ANOVA for grouped tables. We used Bonferroni correction to adjust for multiple comparisons as indicated and used Dunnett T3 correction in one-way ANOVA assuming different standard deviations. For non-normal data distribution, we used Mann-Whitney test for two columns and Kruskal-Wallis test for more than two columns. For podometrics, we analyzed data with Minitab Statistical Software v19 (Minitab 17 Statistical Software (2010), Minitab, State College, PA). We assessed differences between two groups with unpaired, two-sample Student’s t-test. We adjusted by Welch’s correction for unequal variances when necessary. For RNAseq analysis, we compared DESeq2-normalized, log-transformed gene counts (log2(normalized counts + 1)) for comparisons as indicated, and used the q-value function from q-value R package with default settings to run adjustment for multiple comparisons. We used the online software Metascape (http://metascape.org/) tool to perform gene ontology (GO) analysis ^96^ using genes with q-values less than 0.05 as inputs. We ran this analysis with default parameters to infer a hierarchical clusterization of significant ontology terms into a tree based on Kappa-statistical similarities among gene memberships. We selected the term with the best q-value within each cluster as its representative term. We retained ontology clusters with a q-value of less than 0.05 as significantly enriched. We designated statistical significance at a 2-sided p-value < 0.05 for all biologically meaningful comparisons. For each experiment we note the number of samples in the corresponding figure legend. We represent data per mouse when indicated and include mean ± SEM.

### Study Approval

The Palo Alto Medical Foundation Institutional Review Board (IRB) approved the collection and storage of human samples and de-identified clinical data. The Stanford University School of Medicine Institutional Review Board (IRB) approved the collection and storage of human samples and de-identified clinical data from the Stanford Precision Biobank. The Stanford Research Compliance Office Administrative Panel for the Protection of Human Subjects – IRB approved the use of stored human sera. The Stanford Institutional Animal Care and Use Committee approved the experiments in mice.

## Abbreviations

DKD: diabetic kidney disease
Esm-1: endothelial cell-specific molecule-1

## Author Contributions

XZ, LEH, ML, PS, AR, JSM, NFC, and VB conceived of experiments. XZ, AG, AB, CL, PaN, XY, BS, BG, XXW, and KM performed experiments. XZ, LEH, AG, MS, LP, BS, XW, PrN, AR, and JSM analyzed data. XZ and VB drafted the manuscript. XZ, LEH, AG, LP, ML, AR, JSM, NFC, and VB edited the manuscript. All authors approved of the final manuscript.

## Acknowledgements

pCMV(CAT)T7-SB100 was a gift from Zsuzsanna Izsvak (Addgene plasmid # 34879).

Emory University Integrated Genomics Core subcloned mouse Esm-1 over-expression plasmids. The Stanford Shared FACS Facility provided flow cytometer, training, and sorting services. The Stanford Animal Histology Service provided services for paraffin embedding and sectioning. The Stanford Canary Center Imaging Facility provided the Nanozoomer Digital Scanner. This work utilized computing resources and differential gene expression analysis provided by the Stanford Genome Center. The authors thank Drs. Alan Pao, Glenn Chertow, and Philippe Lassalle for helpful discussions and critical review of the manuscript. XZ was funded, in part, by a Larry L. Hillblom Foundation Postdoctoral Fellowship award and the Holmgren Family Foundation. LEH was funded by two NIH training grants (T32 DK007357, T32 AI07290) and an American Heart Association / Enduring Hearts postdoctoral fellowship (17POST33660597). XY and AR were funded by R01 DK116567. XXW, KM, and ML were funded by R01 DK116567 and NIH R01 DK127830. PS was funded by R01 DK114485. JSM was funded by a Veteran’s Administration Merit Award (I01 CX001971). Financial support for this work was also provided, in part, by the NIDDK Diabetic Complications Consortium (DiaComp, www.diacomp.org, DK076169). VB was funded, in part, by the NIDDK (R01 DK091565).

## Notes

**Conflicts of Interest** JSM has a family member who is employed by and has an equity interest in Genentech/Roche. NFC receives salary support, in part, from Biothelis. VB has a family member who receives speaking fees from CareDx.

https://www.ncbi.nlm.nih.gov/geo/query/acc.cgi?acc=GSE175449

### Competing Interest Statement

JSM has a family member who is employed by and has an equity interest in Genentech/Roche. NFC receives salary support, in part, from Biothelis. VB has a family member who receives speaking fees from CareDx.

