## Supplementary Materials for "Endothelial Cell-Specific Molecule-1 Inhibits Albuminuria in Diabetic Mice"

#### Supplementary Figures

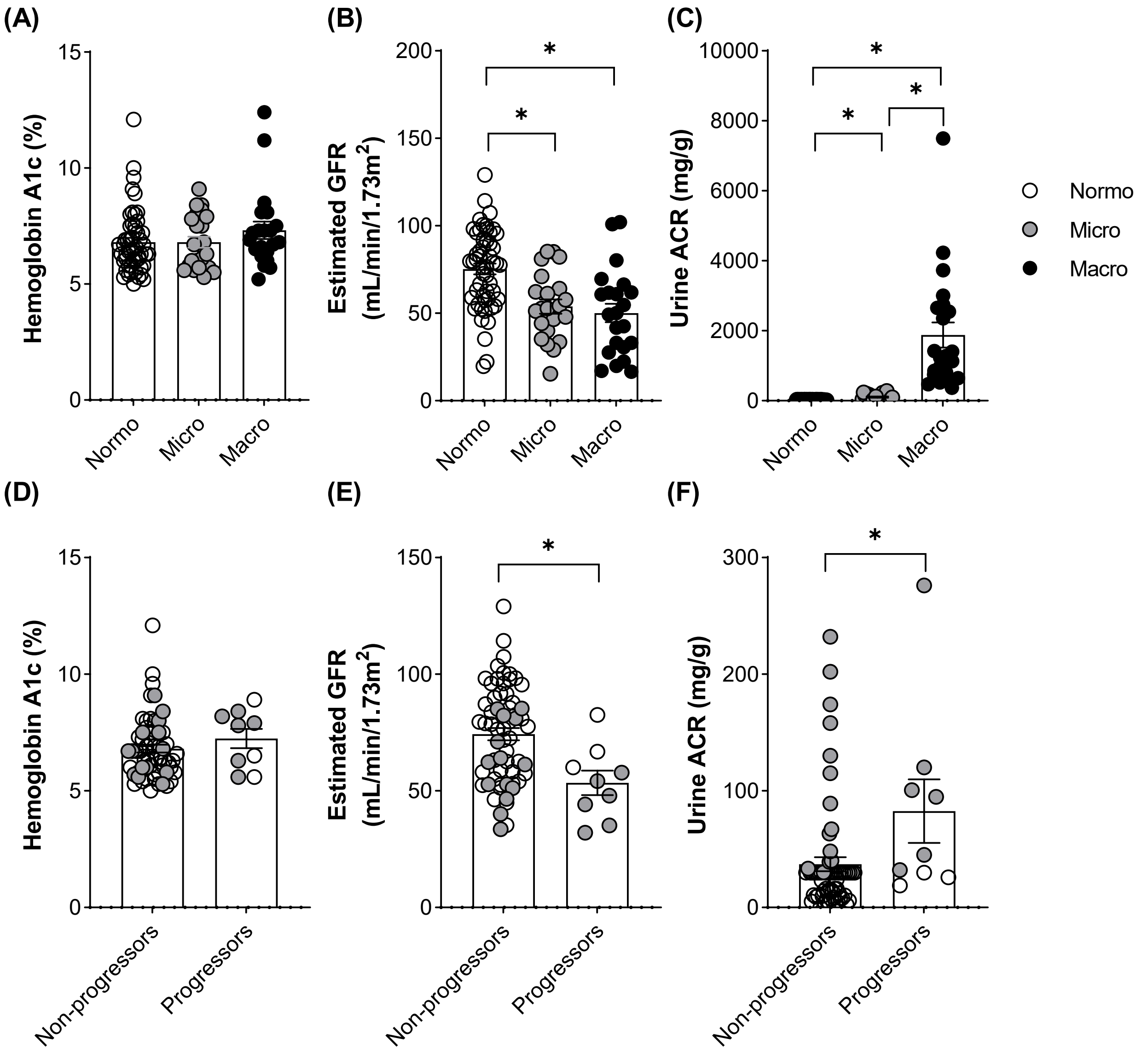

**Supplementary Figure 1: Characteristics of patients with diabetes.** (A) Hemoglobin A1c, (B) estimated glomerular filtration rate, and (C) urine albumin-to-creatinine ratio (ACR) for patients classified by degree of albuminuria. * $p < 0.05$ by one-way ANOVA. N= 21-54 patients per group. (D) Hemoglobin A1c, (E) estimated glomerular filtration rate, and (F) urine albumin-to-creatinine ratio (ACR) for patients classified as non-progressors vs. progressors. N= 9-66 patients per group. White, patients with diabetes with normoalbuminuria; Grey, patients with microalbuminuria; and Black, patients with macroalbuminuria. * $p < 0.05$ compared across groups by unpaired Student's t-test.

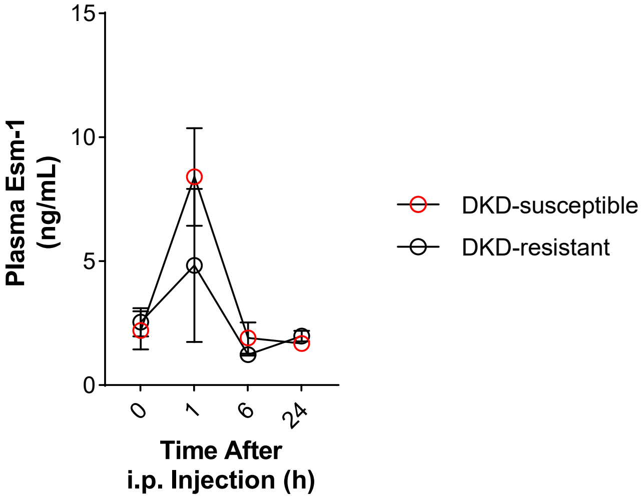

**Supplementary Figure 2: Intraperitoneal injection of recombinant protein only transiently increases circulating mouse Esm-1.** We injected DKD-susceptible and DKD-resistant mice intraperitoneally (i.p.) with recombinant mouse Esm-1 and quantified plasma Esm-1 at indicated time points. *Red*, DKD-susceptible mice; *Black*, DKD-resistant mice. Results are presented as mean ± SEM. N= 2 mice per group.

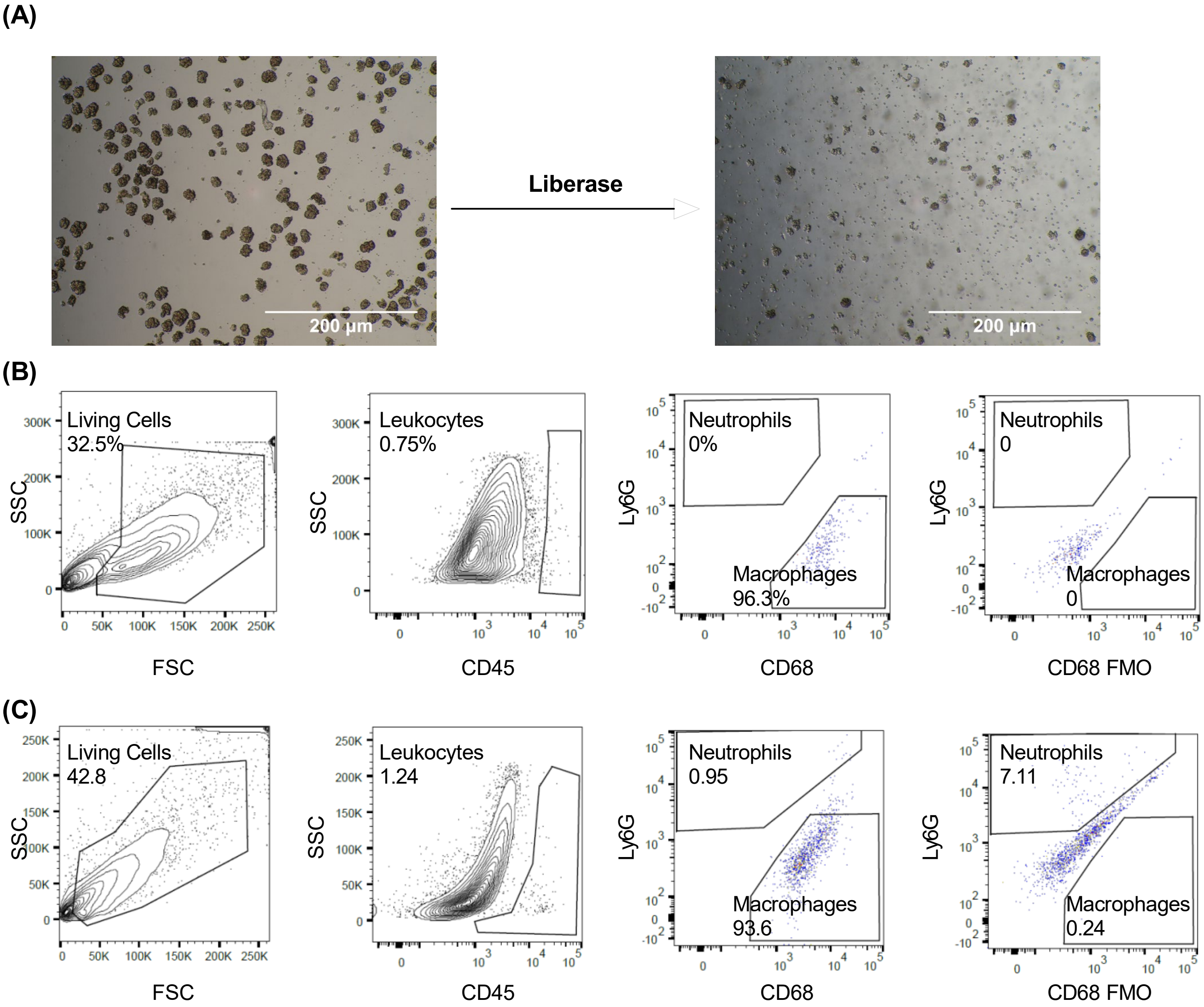

**Supplementary Figure 3: Method of glomerular leukocyte infiltration.** (A) Brightfield images of glomeruli extracted by collagenase digestion and sieving, followed by 200 µg/mL liberase for 24 hours. Scale bars (white) as indicated. Representative gating of (B) glomerular and (C) tubulointerstitial cells using flow cytometry. First column, gating for living cells selected from unity axis of forward scatter (FSC) and side scatter (SSC) as indicated; Second column, gating for CD45(+) cells (i.e. leukocytes) from living cells by side scatter (SSC) and CD45-Brilliant Violet 421 as indicated; Third column, gating for CD68(+)Ly6G(-) cells (i.e. macrophages) and CD68(-)Ly6G(+) cells (i.e. neutrophils) from CD45(+) cells by Ly6G-PE and CD68-Alexa Fluor 647 as indicated; Fourth column, gating windows determined by fluorescence minus one (FMO) controls as indicated.

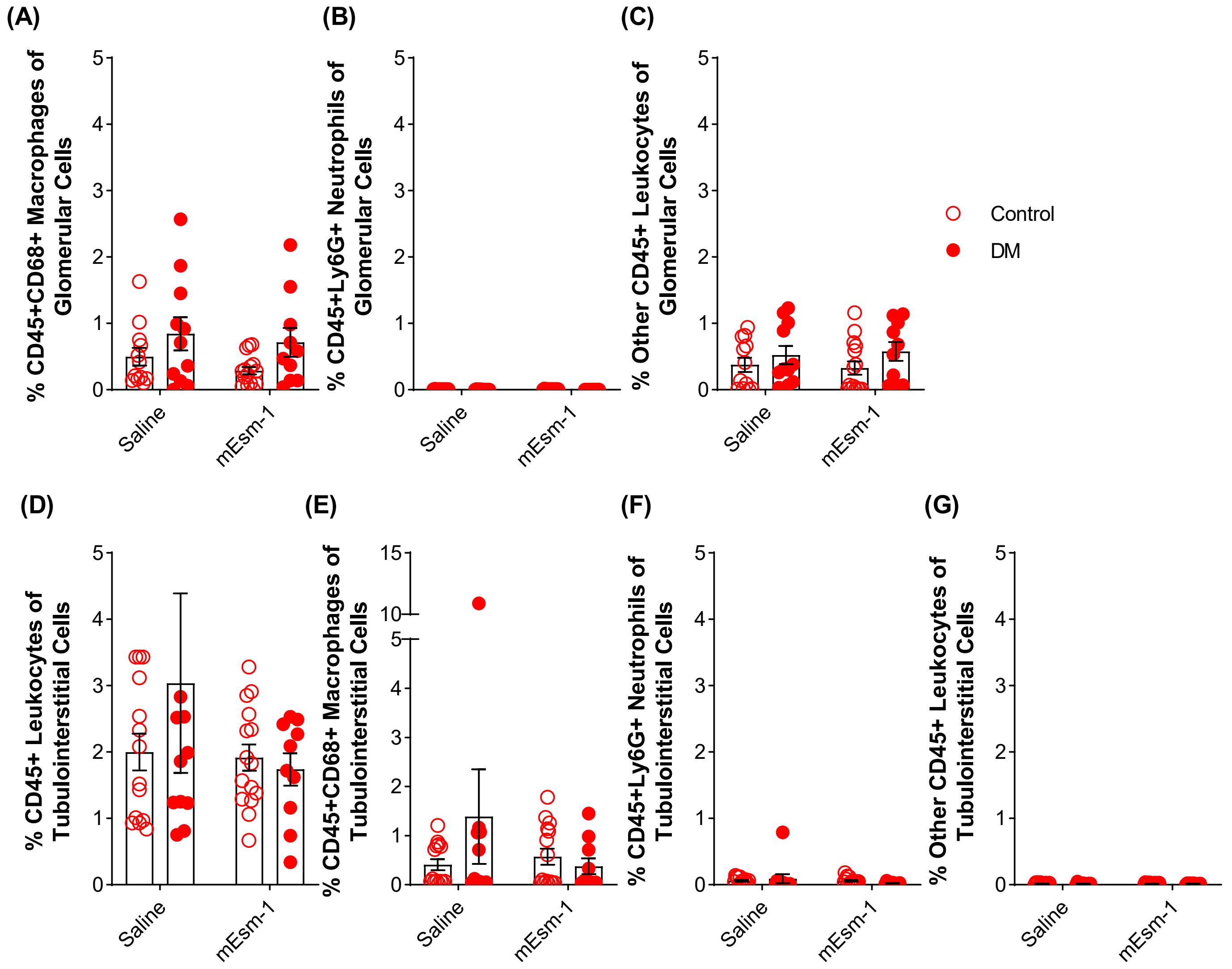

**Supplementary Figure 4: Systemic over-expression of mouse Esm-1 does not alter infiltration of leukocyte subsets in glomerular or tubulointerstitial compartments in diabetic DKD-susceptible mice.** Flow cytometry analysis from glomeruli and tubulointerstitium of control and diabetic mice treated with saline or mouse Esm-1-injection as indicated. Percentage of (A) glomerular macrophages, (B) glomerular neutrophils, (C) other glomerular leukocyte cell types, (D) tubulointerstitial leukocytes, (E) tubulointerstitial macrophages, (F) tubulointerstitial neutrophils, and (G) other tubulointerstitial leukocyte cell types. Open circles, control mice; closed circles, diabetic (DM) mice. Results are presented per mouse and include mean ± SEM. * $p < 0.05$ compared across groups as indicated; # $p < 0.05$ vs. control mice by two-way ANOVA. N= 10-15 mice per group.

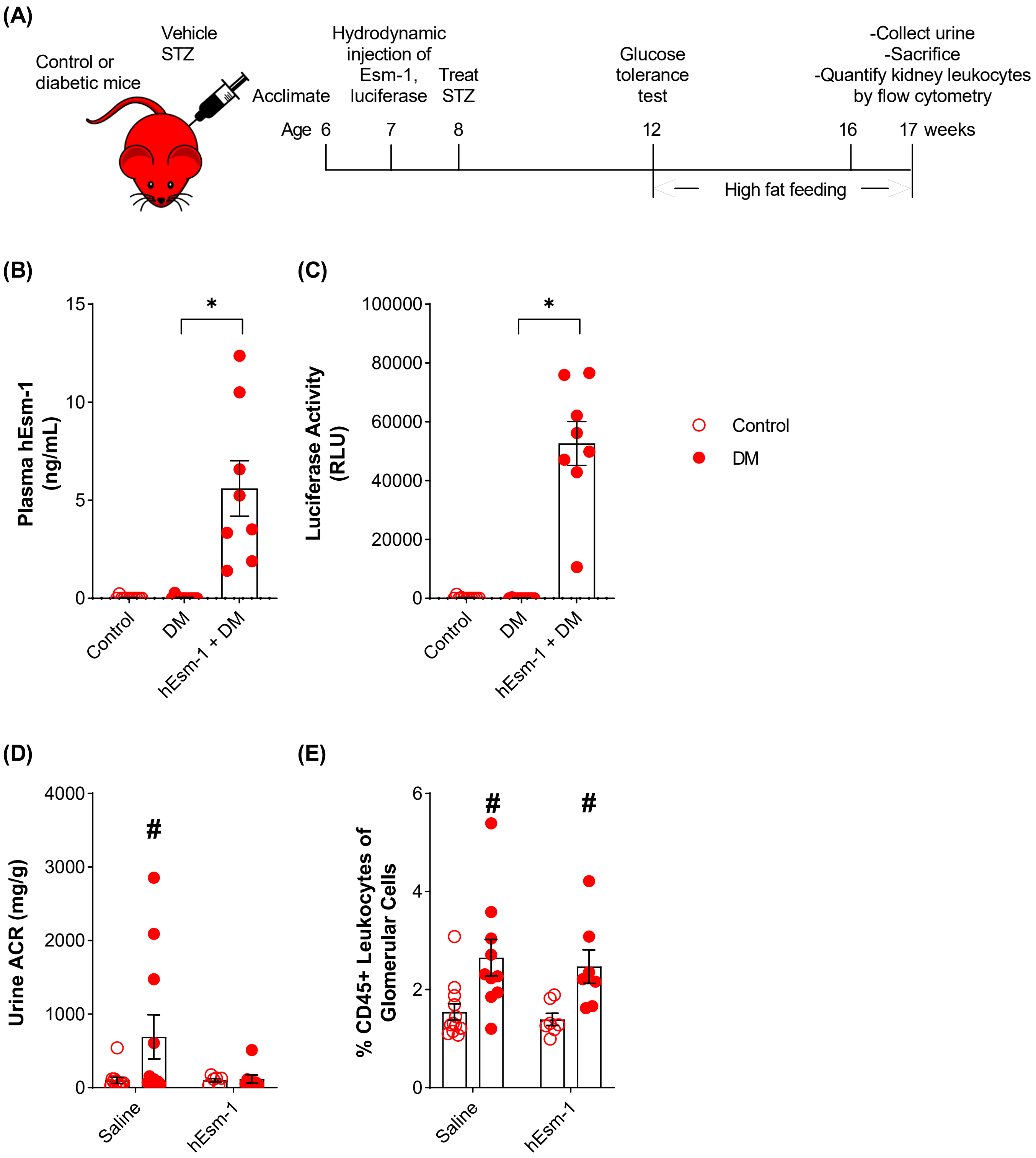

**Supplementary Figure 5: Systemic over-expression of human Esm-1 reduces diabetes-induced albuminuria in DKD-susceptible mice.** (A) Timeline of experimental design. We induced expression of human Esm-1 (hEsm-1) by hydrodynamic injection and treated mice with vehicle vs. streptozotocin (STZ) and high fat feeding at indicated time points, followed by sacrifice at 17 weeks of age. We quantified (B) plasma human Esm-1, (C) luciferase activity, (D) urine albumin-to-creatinine ratio (ACR), (E) glomerular leukocytes in control and diabetic mice. Open circles, control mice; closed circles, diabetic (DM) mice. Results are presented per mouse and include mean ± SEM. * p-value < 0.05 compared across as indicated;

### p-value < 0.05 vs.  control mice, by one-way ANOVA (B-C) or two-way ANOVA (D-E). RLU, relative luciferase units. N= 6-12 mice per group.

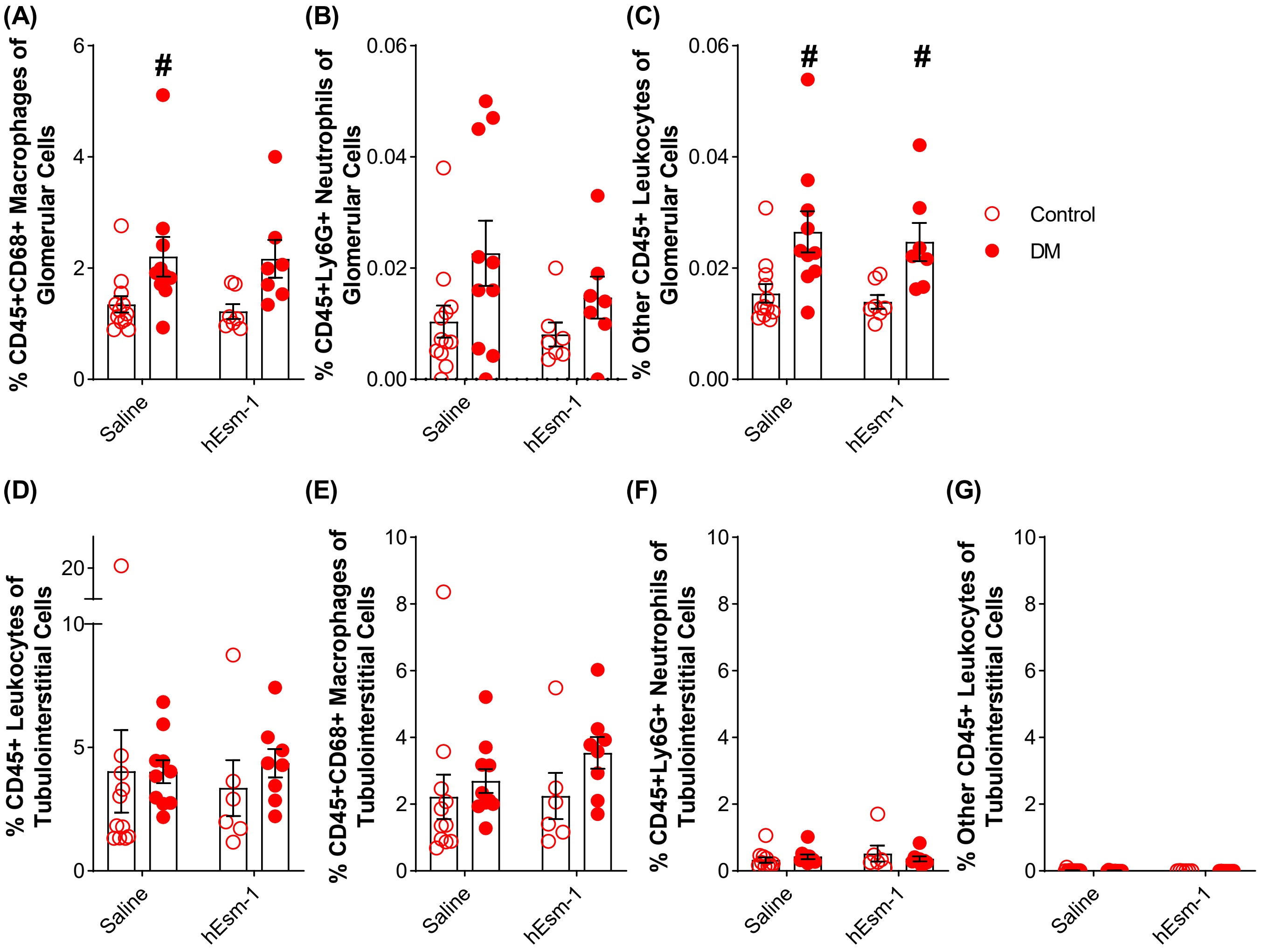

**Supplementary Figure 6: Systemic over-expression of human Esm-1 does not alter infiltration of leukocyte subsets in glomerular or tubulointerstitial compartments in diabetic DKD-susceptible mice.** Flow cytometry analysis from glomeruli and tubulointerstitium of control and diabetic mice treated with saline or mouse Esm-1-injection as indicated. Percentage of (A) glomerular macrophages, (B) glomerular neutrophils, (C) other glomerular leukocyte cell types, (D) tubulointerstitial leukocytes, (E) tubulointerstitial macrophages, (F) tubulointerstitial neutrophils, and (G) other tubulointerstitial leukocyte cell types. Open circles, control mice; closed circles, diabetic (DM) mice. Results are presented per mouse and include mean ± SEM. * $p < 0.05$ compared across groups as indicated; # $p < 0.05$ vs. control mice by two-way ANOVA. N= 6-12 mice per group.

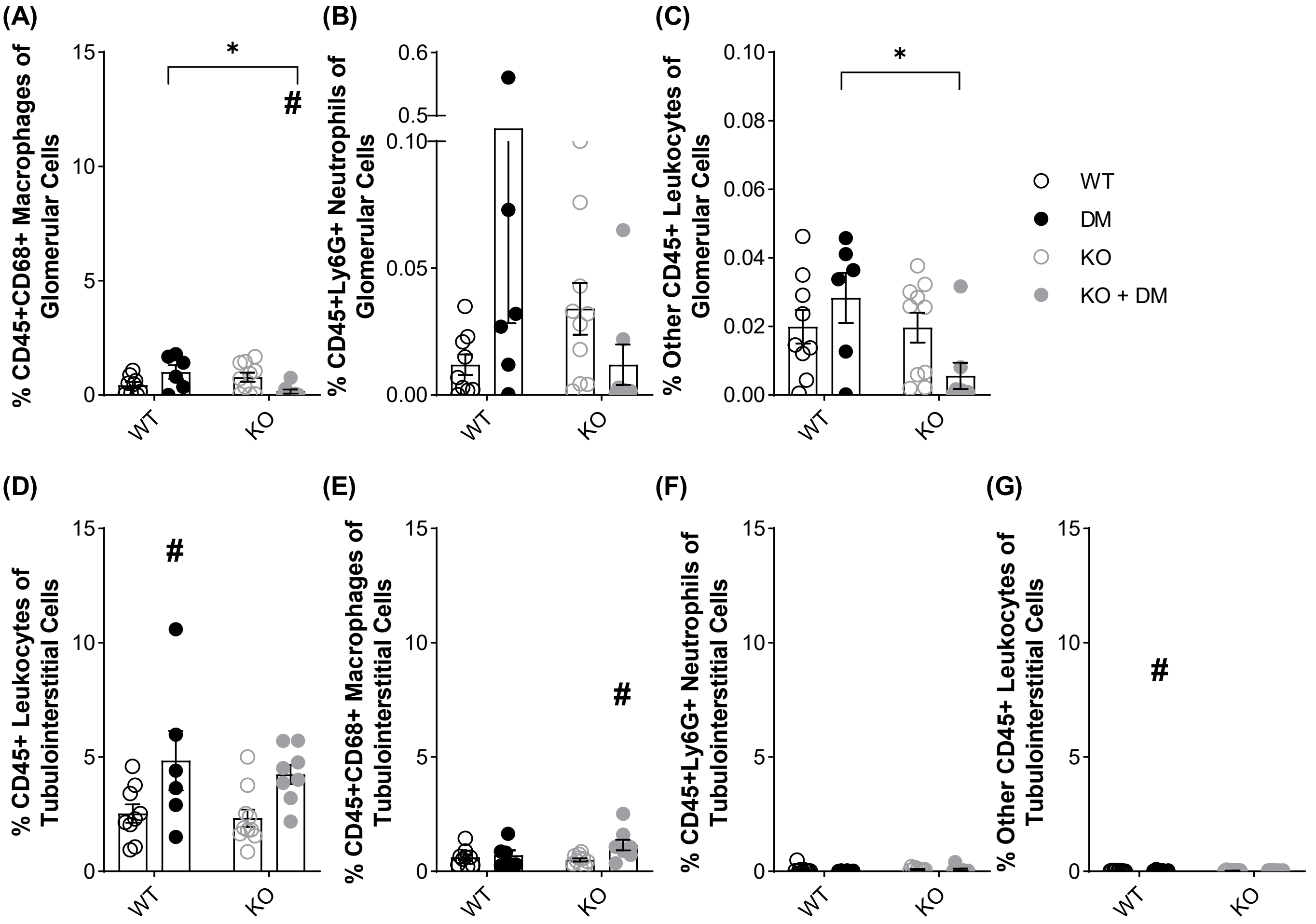

**Supplementary Figure 7: Constitutive deletion of Esm-1 and leukocyte subsets in glomerular and tubulointerstitial compartments in diabetic DKD-susceptible mice.** Flow cytometry analysis from glomeruli and tubulointerstitium of control and diabetic, wild-type and knockout mice treated with saline or mouse Esm-1-injection as indicated. Percentage of (A) glomerular macrophages, (B) glomerular neutrophils, (C) other glomerular leukocyte cell types, (D) tubulointerstitial leukocytes, (E) tubulointerstitial macrophages, (F) tubulointerstitial neutrophils, and (G) other tubulointerstitial leukocyte cell types. Open circles, control mice; closed circles, diabetic (DM) mice. *Black*, wild-type (WT) littermate controls. *Grey*, Esm-1 knockout (KO) mice. are presented per mouse and include mean ± SEM. * $p < 0.05$ compared across groups as indicated; # $p < 0.05$ vs. control mice by two-way ANOVA. N= 6-10 mice per group.

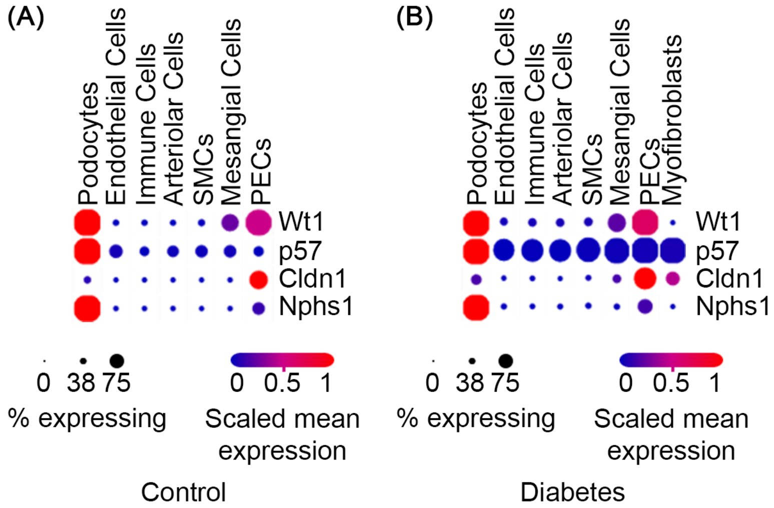

**Supplementary Figure 8: Parietal and visceral podocyte markers in mouse kidney.** Scaled mean expression and percentage of expressing cells for Wt1, p57, Cldn1, and Nphs2 genes in each glomerular cell type in: (A) control (*ob/het*) and (B) diabetic (*ob/ob*) mice. Based on Cldn1 and Nphs1, podocytes correspond to visceral epithelial cells or visceral podocytes. PECs correspond to parietal epithelial cells or parietal podocytes. *SMC,* smooth muscle cells.

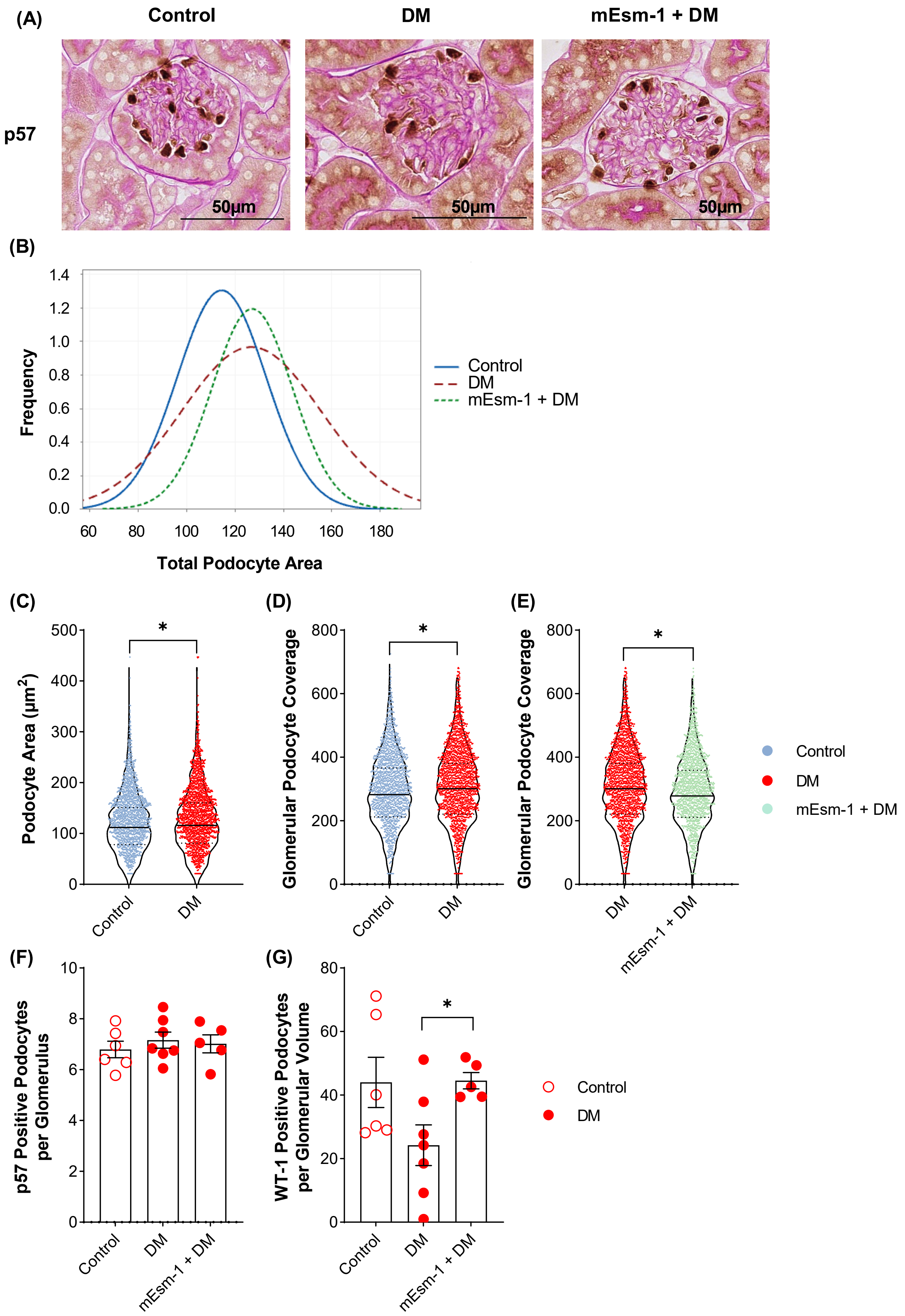

(A)
Control
DM
mEsm-1 + DM
p57
50µm
(B)
Frequency
Total Podocyte Area
Control
DM
mEsm-1 + DM
(C)
Podocyte Area (µm²)
Control
DM
(D)
Glomerular Podocyte Coverage
Control
DM
(E)
Glomerular Podocyte Coverage
DM
mEsm-1 + DM
Control
DM
mEsm-1 + DM
(F)
p57 Positive Podocytes per Glomerulus
Control
DM
mEsm-1 + DM
(G)
WT-1 Positive Podocytes per Glomerular Volume
Control
DM
mEsm-1 + DM
Control
DM

.

**Supplementary Figure 9: Systemic over-expression of mouse Esm-1 attenuates podocyte injury induced by diabetes.** (A) Representative images of mouse kidney sections stained for p57 expression. Scale bars (*black*) as indicated. (B) Histogram of total podocyte nuclear area. N= 5-7 mice per group. (C-G) Analysis of podocytes from mice in (B). Violin plots of (C) podocyte nuclear area, (D,E) glomerular podocyte coverage. *Blue*, control mice; *Red*, diabetic (DM) mice; and *Green*, diabetic mice with mEsm-1 over-expression (mEsm-1 + DM). * $p < 0.05$ by unpaired Student`s t-test. (F) p57(+) podocytes per glomerulus. (G) WT-1(+) podocytes per glomerular volume in the same samples as (F). N=5-7 mice per group. Open circles, control mice; closed circles, diabetic (DM) mice. * $p < 0.05$ compared across groups as indicated by one-way ANOVA.

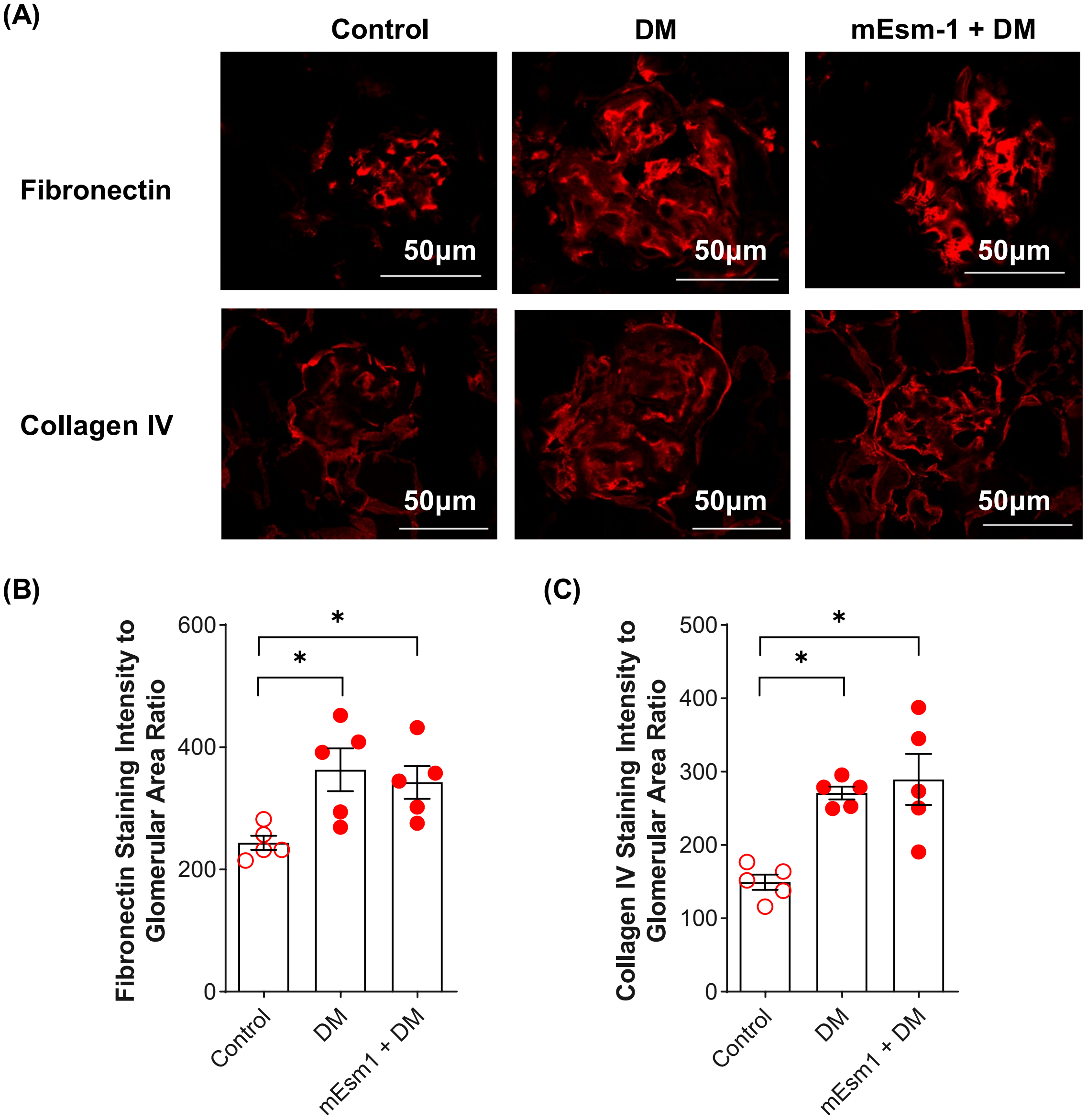

**Supplementary Figure 10: Systemic over-expression of mouse Esm-1 does not influence glomerular extracellular matrix deposition induced by diabetes.** (A) Representative images of mouse kidney sections stained for fibronectin staining or collagen IV. Scale bars (white) as indicated. (B) Glomerular fibronectin and (C) Glomerular collagen IV staining intensity. Open circles, control mice; closed circles, diabetic (DM) mice. Results are presented per mouse and include mean ± SEM. * $p < 0.05$ compared across groups as indicated by one-way ANOVA. N= 5 mice per group.

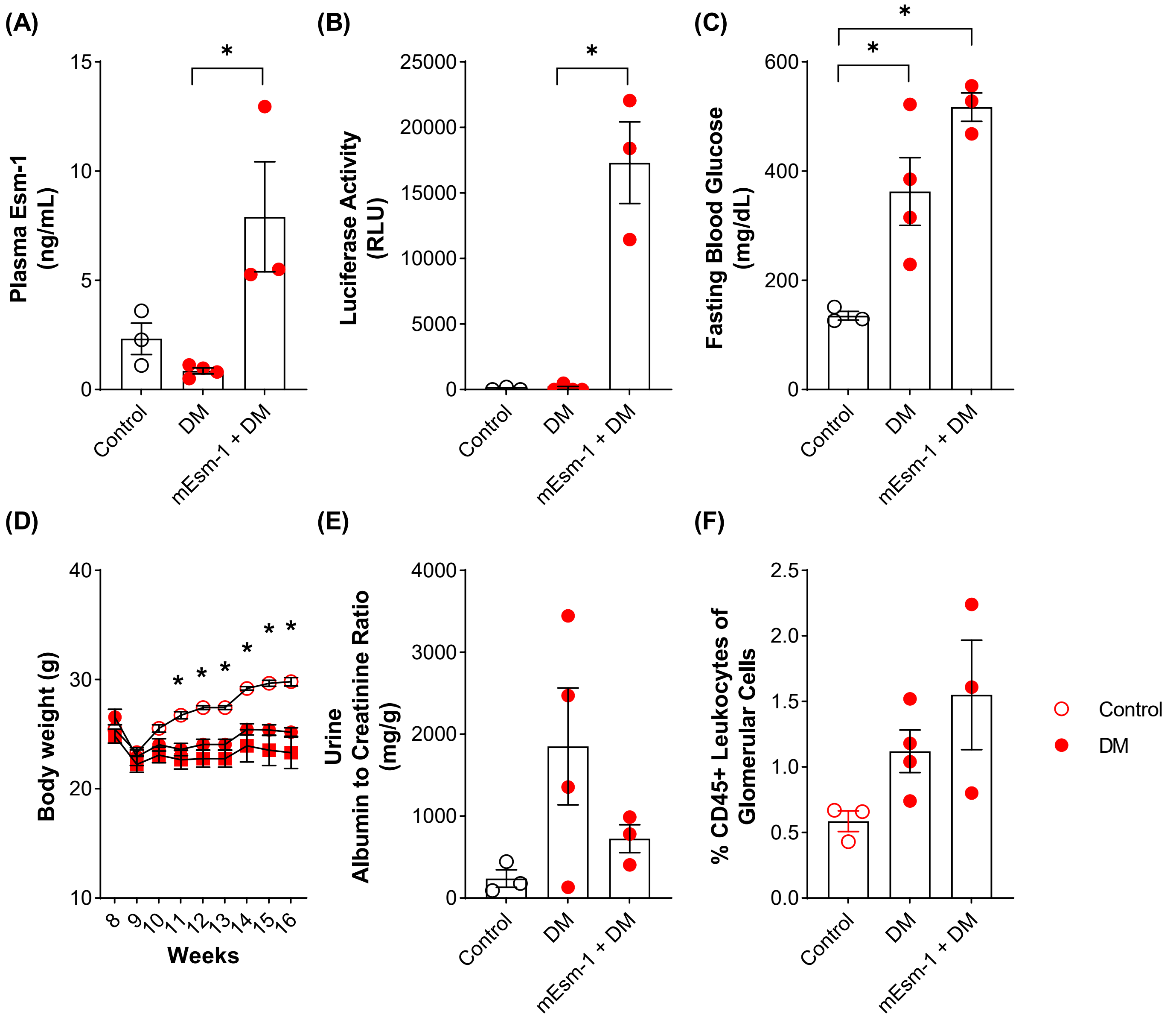

**Supplementary Figure 11: Clinical and histologic characteristics of mice utilized for glomerular RNAseq.** We quantified (A) plasma Esm-1, (B) luciferase activity, (C) fasting blood glucose, (D) body weight, (E) urine albumin-to-creatinine ratio (ACR), and (F) glomerular leukocytes in control and diabetic mice. Open circles, control mice; closed circles, diabetic (DM) mice. In all panels except (D), results are presented per mouse and include mean ± SEM. In (D), values for mice with over-expression of mEsm-1 shown as squares and results are presented as mean ± SEM. RLU, relative luciferase units. * p-value < 0.05 compared across groups as indicated by one-way ANOVA. N= 9-15 mice per group.

DE significant genes (DM vs. Control)

| Gene | Response to interferon-beta | Response to virus | Response to cytokine stimulus | Regulation of innate immune response | Response to type 1 interferon | Response to interferon-alpha | Response to interferon-gamma | $Log_2FC$ | p-value | q-value |
|---|---|---|---|---|---|---|---|---|---|---|
| AI607873 | ● | | | | | | | 2.52323041 | 5.70E-12 | 2.37E-09 |
| Bst2 | ● | ● | | | | ● | ● | 1.66907319 | 7.21E-07 | 0.0001143 |
| Gbp2 | ● | ● | | | | | ● | 1.70783557 | 3.49E-23 | 5.28E-20 |
| Gbp3 | ● | ● | | | | | ● | 2.19000693 | 3.25E-22 | 4.16E-19 |
| Gbp6 | ● | ● | | | | | ● | 2.86500555 | 7.07E-18 | 5.88E-15 |
| Gm4951 | ● | | | | | | ● | 1.52061603 | 1.89E-05 | 0.00186148 |
| Ifi202b | ● | | | | | | | 2.55892785 | 1.92E-15 | 1.28E-12 |
| Ifi203 | ● | | | ● | | | | 1.30596001 | 1.13E-07 | 2.23E-05 |
| Ifi204 | ● | | | ● | | ● | | 2.33262472 | 3.16E-18 | 3.09E-15 |
| Ifi47 | ● | | | | | | ● | 2.45976907 | 1.40E-07 | 2.56E-05 |
| Ifit1 | ● | ● | | | | ● | | 2.5702461 | 2.32E-24 | 4.83E-21 |
| Ifit3 | ● | ● | | | | ● | | 2.80456444 | 1.11E-35 | 6.16E-32 |
| Ifitm3 | ● | ● | ● | | ● | ● | ● | 1.00177899 | 3.80E-06 | 0.00051009 |
| Igtp | ● | | | | | | ● | 1.54257549 | 2.82E-08 | 6.52E-06 |
| Iigp1 | ● | ● | ● | | | | ● | 1.65545969 | 4.22E-06 | 0.00055748 |
| Irgm1 | ● | ● | ● | ● | | | ● | 1.17629976 | 2.89E-08 | 6.59E-06 |
| Irgm2 | ● | ● | ● | ● | | | ● | 1.0431233 | 0.00010487 | 0.00715408 |
| Mnda | ● | | | ● | | | | 1.99132069 | 8.12E-06 | 0.00091944 |
| Mndal | ● | | | ● | | | | 2.65129115 | 0.00040164 | 0.02102269 |
| Ndufa13 | ● | | | | | | | 0.95135892 | 9.35E-05 | 0.00653911 |
| Pydc3 | ● | | | ● | | | | 3.45520008 | 0.00068458 | 0.0325488 |
| Stat1 | ● | ● | ● | ● | ● | | ● | 0.87985149 | 3.41E-05 | 0.00293635 |
| Tgtp2 | ● | | | | | | ● | 1.44391957 | 0.00022522 | 0.01338829 |
| Xaf1 | ● | | | | | | | 1.7004419 | 1.40E-14 | 7.77E-12 |
| Apoe | | ● | | ● | | | | 1.06689293 | 5.26E-06 | 0.0006658 |
| B2m | | ● | | ● | | | ● | 1.30084798 | 1.37E-10 | 4.96E-08 |
| Batf2 | | ● | | | | | ● | 1.71652208 | 4.77E-06 | 0.00061935 |
| Bcl2l11 | | ● | | | | | | 0.92368584 | 0.00020912 | 0.01261154 |
| Cxcl10 | | ● | ● | | | | | 3.31401266 | 6.05E-15 | 3.60E-12 |
| Cxcl13 | | ● | ● | | | | ● | 3.5848997 | 5.52E-08 | 1.16E-05 |
| Cxcl2 | | ● | ● | | | | | 0.75477174 | 0.00023565 | 0.01386029 |
| Cxcl9 | | ● | ● | | | | | 4.56410913 | 0.00080938 | 0.03656722 |
| Cxcr4 | | ● | ● | | | | | 1.71072986 | 0.00028239 | 0.01615223 |
| Ddx58 | | ● | ● | ● | | | | 1.38829624 | 7.43E-09 | 2.03E-06 |
| Ddx60 | | ● | | ● | | | | 3.69925052 | 4.72E-07 | 7.70E-05 |
| Denr | | ● | | | | | | 0.94749535 | 0.00064503 | 0.03085218 |
| Dhx58 | | ● | | ● | | | | 2.83720155 | 3.87E-08 | 8.48E-06 |
| Dtx3l | | ● | | ● | | | | 1.01927458 | 3.31E-05 | 0.00289327 |
| Eif2ak2 | | ● | | ● | | ● | | 1.36728233 | 2.28E-08 | 5.63E-06 |
| Gbp7 | | ● | | | | | ● | 1.49220684 | 6.82E-14 | 3.55E-11 |
| Gbp8 | | ● | | | | | ● | 3.6198876 | 5.66E-25 | 1.57E-21 |
| Grn | | ● | | ● | | | ● | 0.71060702 | 0.00098419 | 0.04274163 |
| H2-K1 | | ● | | ● | | | ● | 1.20650086 | 2.11E-13 | 1.06E-10 |
| H2-T23 | | ● | | ● | | | ● | 1.41930493 | 5.33E-11 | 2.02E-08 |
| I830012O16Rik | | ● | | | | | | 3.31108647 | 1.21E-24 | 2.88E-21 |
| Ifi27l2a | | ● | | | | | | 5.58560694 | 8.49E-10 | 2.77E-07 |
| Ifih1 | | ● | ● | ● | | | | 2.37323174 | 2.28E-17 | 1.73E-14 |
| Ifit2 | | ● | | | | ● | | 3.32790823 | 3.63E-44 | 3.02E-40 |
| Il6 | | ● | ● | ● | | | ● | 1.33695286 | 4.15E-12 | 1.82E-09 |
| Inhbb | | ● | | ● | | | | 3.25295278 | 3.41E-18 | 3.15E-15 |
| Irf7 | | ● | ● | ● | ● | | | 2.64796086 | 1.22E-28 | 4.06E-25 |
| Isg15 | | ● | | ● | ● | | ● | 1.72397606 | 3.40E-09 | 1.01E-06 |
| Isg20 | | ● | | | | | | 3.63429129 | 2.98E-09 | 9.19E-07 |
| Itgb3 | | ● | | | | | | 1.23438888 | 1.29E-07 | 2.41E-05 |
| Lgals9 | | ● | | ● | | | ● | 1.1790041 | 9.91E-05 | 0.00683683 |
| Mx1 | | ● | | | | | ● | 4.95170883 | 3.34E-06 | 0.00046718 |
| Mx2 | | ● | | | | | ● | 2.14835698 | 4.58E-08 | 9.90E-06 |
| Oas1a | | ● | | | | | | 2.13517369 | 1.93E-06 | 0.0002818 |
| Oas1g | | ● | | | | | | 3.7169368 | 0.0002594 | 0.01509661 |
| Oas2 | | ● | ● | | ● | | | 4.13733148 | 1.30E-15 | 9.02E-13 |
| Oasl1 | | ● | | | | | | 2.19159991 | 6.36E-18 | 5.57E-15 |
| Oasl2 | | ● | | | | | | 2.04740264 | 6.42E-15 | 3.68E-12 |
| Papss1 | | ● | | | | | | 0.9137657 | 1.50E-05 | 0.00150407 |
| Parp10 | | ● | | | | | | 1.03253252 | 0.00014568 | 0.00936207 |
| Parp9 | | ● | ● | ● | | | ● | 1.13502543 | 4.24E-05 | 0.0034938 |
| Pml | | ● | | ● | | | | 0.96775075 | 1.91E-05 | 0.00187012 |
| Rsad2 | | ● | | ● | | | | 3.44931449 | 2.87E-68 | 4.78E-64 |
| Rtp4 | | ● | | | | | | 1.6991708 | 6.57E-11 | 2.43E-08 |
| Serinc3 | | ● | | | | | | 0.48762442 | 0.0004749 | 0.02417348 |
| Serpine1 | | ● | | ● | | | ● | 0.68400121 | 5.30E-06 | 0.0006658 |
| Slfn9 | | ● | | | | | | 5.47524396 | 5.58E-06 | 0.00069313 |
| Tlr2 | | ● | ● | ● | | | ● | 1.4668926 | 0.00016545 | 0.01047091 |
| Tlr3 | | ● | | ● | | | | 1.77747074 | 0.00035427 | 0.01939734 |
| Trim26 | | ● | | | | | | 0.99234183 | 3.25E-05 | 0.00287104 |
| Trim30a | | ● | | ● | | | | 2.55835757 | 3.10E-23 | 5.16E-20 |
| Trim56 | | ● | | ● | ● | | | 0.66664478 | 0.00073106 | 0.03398997 |
| Vgf | | ● | | | | | ● | 1.3711862 | 0.00030861 | 0.01729578 |
| Zbp1 | | ● | ● | ● | ● | | | 3.90786161 | 3.59E-08 | 7.97E-06 |
| Aes | | | ● | | | | | 0.6635171 | 0.00010772 | 0.0072299 |
| Cxcl11 | | | ● | | | | | 3.98013955 | 2.80E-08 | 6.52E-06 |
| Nlrc5 | | | ● | ● | ● | | ● | 0.99237875 | 1.05E-05 | 0.00112034 |
| Parp14 | | | ● | ● | | | ● | 0.98732674 | 6.55E-14 | 3.52E-11 |
| Wnk1 | | | ● | ● | | | | 0.49820844 | 3.32E-05 | 0.00289327 |
| Bsg | | | | ● | | | | 0.54595125 | 0.00033052 | 0.01839991 |
| C4b | | | | ● | | | | 2.30789815 | 0.00044634 | 0.02292987 |
| Cd274 | | | | ● | | | | 2.43369776 | 7.89E-32 | 3.28E-28 |
| Cyp1b1 | | | | ● | | | | 0.92437196 | 0.00032471 | 0.01813685 |
| Cyp2j6 | | | | ● | | | | 1.18121921 | 0.00075841 | 0.03496893 |
| Fabp4 | | | | ● | | | | 1.20241343 | 4.11E-05 | 0.00340353 |
| Gbp5 | | | | ● | | | ● | 1.60440039 | 5.59E-13 | 2.74E-10 |
| H2-D1 | | | | ● | | | | 0.92779358 | 1.27E-05 | 0.00128897 |
| H2-Q10 | | | | ● | | | | 1.70358414 | 0.00016866 | 0.01059376 |
| H2-Q4 | | | | ● | | | | 1.08846776 | 4.14E-16 | 3.00E-13 |
| H2-T24 | | | | ● | | | | 1.85215171 | 1.19E-07 | 2.30E-05 |
| Hilpda | | | | ● | | | | 1.19851794 | 0.00023169 | 0.0136754 |
| Ndufc2 | | | | ● | | | | 1.07187395 | 7.98E-05 | 0.00582575 |
| Pigr | | | | ● | | | | 1.47596546 | 4.80E-06 | 0.00061935 |
| Pnp | | | | ● | | | ● | 0.79996955 | 3.92E-05 | 0.00327881 |
| Slc3a2 | | | | ● | | | | -0.455432 | 0.00035944 | 0.01948831 |
| Sorl1 | | | | ● | | | | 0.80836178 | 0.00117013 | 0.04864727 |
| Tap1 | | | | ● | | | | 1.21456067 | 1.90E-06 | 0.00027987 |
| Trafd1 | | | | ● | | | | 1.17882683 | 2.81E-08 | 6.52E-06 |
| Usp18 | | | | ● | | | | 2.98201543 | 2.12E-18 | 2.21E-15 |
| Gem | | | | | | | ● | 0.86707168 | 8.61E-06 | 0.00095542 |
| Pck1 | | | | | | | ● | 0.9765771 | 0.00097246 | 0.04237307 |
| Rab27a | | | | | | | ● | 1.79306898 | 0.00016684 | 0.01051907 |

DE significant genes (mEsm-1 + DM vs. Control)

| Gene | Response to interferon-beta | $Log_2FC$ | p-value | q-value |
|---|---|---|---|---|
| AI607873 | ● | 1.94138 | 6.26E-08 | 0.00000373 |
| Bst2 | ● | 1.56467 | 0.0000026 | 0.0000924 |
| Gbp2 | ● | 1.33237 | 1.56E-12 | 4.04E-10 |
| Gbp3 | ● | 1.96401 | 2.98E-18 | 2.52E-15 |
| Gbp6 | ● | 2.83864 | 2.78E-18 | 2.52E-15 |
| Ifi202b | ● | 2.18635 | 2.01E-12 | 5.06E-10 |
| Ifi203 | ● | 0.97145 | 0.0000817 | 0.00155932 |
| Ifi204 | ● | 2.08455 | 8.29E-15 | 3.86E-12 |
| Ifi47 | ● | 1.86768 | 0.0000683 | 0.00134491 |
| Ifit1 | ● | 2.17433 | 1.19E-18 | 1.23E-15 |
| Ifit3 | ● | 2.3494 | 1.17E-24 | 3.63E-21 |
| Igtp | ● | 0.89885 | 0.00132425 | 0.0144868 |
| Iigp1 | ● | 1.0229 | 0.00271508 | 0.02498789 |
| Irgm1 | ● | 0.71976 | 0.00078444 | 0.00964126 |
| Mnda | ● | 1.94875 | 0.00000363 | 0.00012108 |
| Ndufa13 | ● | 1.3005 | 0.00000004 | 0.00000257 |
| Pydc3 | ● | 3.17447 | 0.00154581 | 0.01619519 |
| Stat1 | ● | 0.62109 | 0.00428392 | 0.0354352 |
| Xaf1 | ● | 1.41637 | 4.64E-10 | 6.45E-08 |

**Supplementary Figure 12: Esm-1 attenuates diabetes-induced interferon signaling in glomeruli.** List of interferon-related genes in significantly up-regulated interferon pathways in Figure 7A and corresponding $log_2$ fold change, p-value, and q-value in comparison of diabetes (DM) vs. control and diabetes + mouse Esm-1 (mEsm-1 + DM) vs. control. *Black*, genes significantly regulated with diabetes vs. control, independent of Esm-1 over-expression; *Red*, genes that are not significantly regulated with Esm-1 treatment.

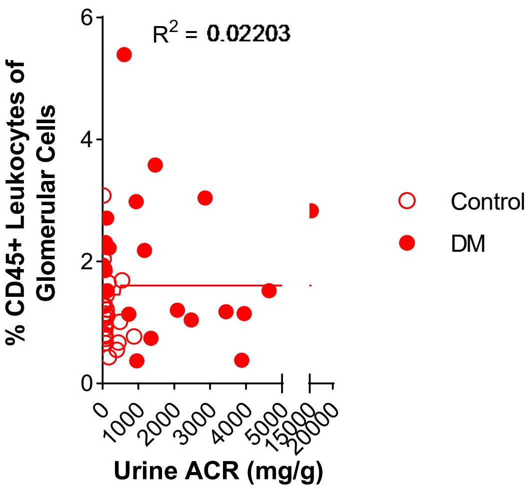

**Supplementary Figure 13: Correlation of glomerular leukocyte infiltration and albuminuria in diabetic DKD-susceptible mice.** Open circles, control mice; closed circles, diabetic (DM) mice. Results are presented per mouse. $R^2$ for non-linear regression (solid line) is shown. N= 23-26 mice per group.

**Supplementary Methods**

*Intraperitoneal Injection of Recombinant Esm-1 Protein*

We injected 8-week old male mice intraperitoneally with 1.7 µg mouse recombinant Esm-1 (R&D, Minneapolis, MN). We collected retro-orbital blood at the indicated time points and measured plasma Esm-1 by ELISA (Aviscera Biosciences, Santa Clara, CA), following the manufacturer's instructions.

*Quantification of Glomerular Fibronectin and Collagen IV*

We stained kidney frozen sections for fibronectin (Sigma-Aldrich, St. Louis, MO) and collagen type IV (Rockland Immunochemicals, Limerick, PA) and imaged them with a laser scanning confocal microscope (SP8, Leica Biosystems, Buffalo Grove, IL). We quantified the expression level as the sum of pixel values per glomerular area using NIH ImageJ 1.44 software.

*Single-cell RNAseq of mouse glomeruli*

We obtained glomerular single-cell RNAseq data for control (*ob/het*) and diabetic (*ob/ob* 12 weeks) mice from the Broad Institute portal (https://singlecell.broadinstitute.org/single_cell/study/SCP975/single-cell-rna-seq-of-mouse-glomerulus?genes=Wt1%2CCdkn1c&cluster=ob_ob%20wk12&spatialGroups=-&annotation=Cell_type--group--cluster&subsample=all&tab=dotplot#study-visualize)[1]. We used Wt1, Cdkn1c, Cldn1, and Nphs2 as inputs for WT-1, p57, claudin 1, and podocin, respectively.

*Podocyte Staining and Podometrics*

We followed the protocol outlined by Santo et al.[2]. Briefly, we stained 4µm paraffin embedded mouse kidney sections with anti-p57 antibody (ab75974, Abcam, Cambridge, UK)(a marker of terminal podocyte differentiation[3]), incubated with HRP Unovue Rabbit HRP detection reagent (RU-HRP1000, Diagnostic BioSystems, Pleasanton, CA), and developed by DAB substrate ((BSB0018A, Bio SB, Santa Barbara, CA). We further stained sections with periodic acid (Sigma-Aldrich) and Schiff reagent (Sigma-Aldrich), but not counter stained with hematoxylin. We acquired whole slide bright field images with an Aperio AT2 microscope (Leica Microsystems, Buffalo Grove, IL) or a NanoZoomer S360 slide scanner. We identified podocytes and renal tissue compartments by image segmentation, quantified podocyte number, podocyte nuclear area, and glomerular area in each region of interest. For each glomerulus unit, we estimated podocyte spatial density using three steps: (1) dividing the podocyte count by the glomerular area, (2) summarizing the total area of podocyte nuclei within the glomerulus, and (3) dividing the cumulative podocyte nuclear area by the glomerular area. Glomerular podocyte coverage is the ratio of total podocyte nuclear area: glomerular area.

**Supplementary Tables**

**Supplementary Table 1: Podometric data.**

| Characteristic | DM vs. Control | mEsm-1 + DM vs. DM |
|---|---|---|
| **Number of mice** | 7 vs. 6 | 4 vs. 7 |
| **Number of glomeruli** | 1314 | 1336 |
| **Podocyte Count** | 0.011* | 0.541 |
| **Glomerulus Area** | 0.715 | <0.001*** |
| **Podocytes per unit Glomerular Area** | 0.006** | <0.001*** |
| **Podocyte Nuclear Area** | <0.001*** | 0.607 |
| **Podocyte Nuclei: Glomerulus Area Ratio** | <0.001*** | <0.001*** |

Significance calculated by two-sample unpaired Student's t-test. * p-value < 0.05; ** p-value < 0.01; *** p-value < 0.001 compared across groups as indicated.

**Supplementary Table 2: Antibodies for flow cytometry experiments.**

| Antibody | Company | Catalog Number | Final Dilution |
|---|---|---|---|
| CD45-Alexa488 | BioLegend, San Diego, CA | 103121 | 1:500 |
| CD45-Alexa647 | BioLegend, San Diego, CA | 103123 | 1:1000 |
| CD45-APC | Miltenyi Biotec, San Diego, CA | 130-102-544 | 1:50 |
| CD45-Brilliant Violet 421 | BioLegend, San Diego, CA | 103134 | 1:400 |
| CD45-PE | Miltenyi Biotec, San Diego, CA | 130-117-498 | 1:50 |
| CD68-Alexa488 | Bio-Rad Abd Serotec, Raleigh, NC | MCA1957A88 | 1:16.7 |
| CD68-Alexa647 | Bio-Rad Abd Serotec, Raleigh, NC | MCA1957A647 | 1:10 |
| CD80-Alexa488 | BioLegend, San Diego, CA | 104716 | 1:16.7 |
| Ly6G-APC | Miltenyi Biotec, San Diego, CA | 130-093-140 | 1:10 |
| Ly6G-PE | Miltenyi Biotec, San Diego, CA | 130-102-392 | 1:10 |
| NGAL-Dylight488 | Novus, Centennial, CO | NBP1-05183G | 1:16.7 |

**Supplementary Table 3: Genotyping primer sequences for Esm-1 knockout mice.**

| Primer Name | Primer Sequence |
|---|---|
| **Forward 1f** | ATGTCTATGTCAGTCTCCTC |
| **Reverse 1r** | CTCCAAAATCCAGAAACCC |
| **Reverse 2r** | CTCTTGCCAGCCTCTCTTGT |

**Supplementary Table 4: RT-qPCR primer sequences for validation of bulk RNAseq.**

| Primer Name | Primer Sequence |
|---|---|
| **Mouse *Ackr2* Forward** | ACAGTCACAGGTTCTCCATCCT |
| **Mouse *Ackr2* Reverse** | ACTGATAGTCCCCAAGACCAGA |
| **Mouse *Cxcl11* Forward** | CTGCTCAAGGCTTCCTTATGTT |
| **Mouse *Cxcl11* Reverse** | CCTTTGTCGTTTATGAGCCTTC |
